# Land-use history causes differences in park nighttime cooling capacity and forest structure

**DOI:** 10.1101/2025.02.13.637130

**Authors:** Isabella C. Richmond, Michael A. Paulauskas, Erica Padvaiskas, Laura Carolina Gonzàlez Sinisterra, Kayleigh Hutt-Taylor, Alec L. Robitaille, Carly D. Ziter

## Abstract

Cities are temporally dynamic ecosystems that experience continuous redevelopment over time. Urban parks, which provide critical benefits to resident wellbeing, are developed on different land-use types and contain natural elements that are susceptible to the effects of historical decision making. Thus, understanding the current day functioning of our city’s ecosystems and planning for the equitable distribution of park benefits in the future requires incorporation and understanding of the impacts of historical decision-making. We measured neighbourhood socio-demographic composition, forest structure, and the cooling effect of 33 sites in parks across Montreal, each with a past land-use in one of three classes: agricultural, forested, or industrial. We asked: 1) what are the effects of historical land-use on current park forest structure, diversity, and consequently the capacity to provide cooling? 2) how do surrounding communities differ around parks of each historical land-use type, and what are the implications for equitable access to cooling? We found that forest structure and cooling capacity differed across past land-use type, and forest structure has complex relationships with park cooling capacity. Our results provide evidence of historical environmental injustice impacting current day cooling capacity for marginalized groups. Previously industrial parks had less cooling capacity at night, while simultaneously being surrounded by communities with higher proportions of immigrants, and lower median incomes than Montreal’s average. However, daytime cooling capacity was similar across past land-use type, highlighting the importance of current management decisions to provide a critical ecosystem service, temperature mitigation, regardless of a site’s history. Planting areas of small, dense forest stands with trees >= 5 cm DBH within urban parks can help augment daytime cooling benefits in the city but may hinder nighttime cooling. To provide both nighttime and daytime cooling, a mixed management strategy of park trees is required, where small and large trees are incorporated at different densities. Finally, we find evidence of gentrification surrounding all park sites, emphasizing the complex socioecological dynamics of green infrastructure and the need for community-led greening projects paired with social housing policies.

## 1 Introduction

Urban greening is gaining popularity as a strategy to improve human wellbeing while combatting climate change and biodiversity loss (Haaland and van den Bosch, 2015; Nelson and Bigger, 2022).However, to more fully understand the benefits provided by urban natural ecosystems we must integrate historical conditions. Urban landscapes experience legacy effects, defined as the present-day impacts of past decisions or events (Ossola et al., 2019), which can in turn affect ecosystem functioning for decades to millennia (Cramer et al., 2009; Foster et al., 2003). An important driver of legacy effects in cities is land (re)development, the process of changing one land cover type to another, often multiple times across decades (Raciti et al., 2011; Ziter and Turner, 2018). Throughout the evolution of cities, patches of land are continually maintained, redeveloped, and abandoned, creating a complex history (Alig et al., 2004; Koebel, 1996). For example, many cities in North America were deindustrialized in the mid-20^th^ century, leading to a series of “revitalization” efforts which redeveloped former industrial areas to mixed-use areas with office buildings, hotels, green spaces, and residences (Koebel, 1996). Urban growth in North American cities also often results in expansion at city edges, where agricultural and forested land surrounding cities are developed into urban land-use types, including parks and green spaces (Dallimer et al., 2015). This mix of urban intensification and/or land abandonment in urban cores combined with the expansion of suburbs at city edges means that urbanization results in contemporary green spaces existing in a variety of spaces and incurring a variety of legacy effects (Alig et al., 2004; Dallimer et al., 2015).

Land-use and (re)development patterns in cities are determined by urban planning, a social endeavor with place and period-specific goals, determined by those in power (Pallagst, 2009). Thus, urban planning and land development in post-colonial North America inevitably and intentionally upholds inequity and injustice (Alexander, 1992; Calmore, 1997; Goetz et al., 2020; Powell, 2008; Ture and Hamilton, 1992), ultimately structuring the ecology of our cities (Schell et al., 2020). Environmental injustice is a documented form of inequity across North American cities, and although urban nature is increasingly considered a type of infrastructure all communities deserve access to, decades of evidence demonstrates that urban planning benefits white, rich communities and endangers low-income communities of colour (Gelobter, 1993; Goetz et al., 2020; Maynard, 2017). Marginalized communities are often relegated to neighbourhoods that are poorly designed (Gelobter, 1993), under serviced (Larsen and Gilliland, 2008; Shah et al., 2016), and have environmental hazards that threaten public health (United Church of Christ, 1987; Waldron, 2018). For example, on the island of Montreal urban heat is correlated with the percentage of low-income residents and education level across neighbourhoods (Fan and Sengupta, 2022). Low income and visible minority residents also experience inequity in access to vegetation (Pham et al., 2012), a key factor in combatting urban heat. Further, neighbourhoods surrounding industrial areas were historically inhabited by working class or impoverished immigrants working in the quarries (Héroux and Martin, 2020; Ndiaye, 2004). Today, populations with higher proportions of visible minorities and low-income households are more likely to live near highly congested parks in Montreal, reducing access to park benefits such as heat mitigation (Jepson et al., 2022). More broadly, areas of high social vulnerability in several Canadian cities have lower urban forest resilience (Landry et al., 2020), a key factor determining the longevity and health of urban nature.

Increasing the presence of urban forests and parks is often regarded as universally positive (Roman et al., 2021). However, a concern of urban greening is the potential perpetuation of societal inequities (Kiani et al., 2023; Sax et al., 2022). This is particularly true for parks developed on industrial land in historically marginalized neighbourhoods. Without strong social housing policy, urban greening can lead to gentrification and displacement of residents (Anguelovski, 2016; Anguelovski et al., 2019). In addition, legacies of historical inequity in neighbourhoods through various mechanisms such as environmental racism and presence of industrial lands has been shown to continue to impact key providers of ecosystem services such as current tree canopy and growth (Healy et al., 2022; Nix et al., 2023). The influence of historical land uses and inequity on current distribution of tree canopy has other important implications which have not yet been thoroughly studied, notably the distribution of urban heat.

Urban trees and forests are a critical component to consider in understanding the link between historical legacies and the benefits of green spaces. Park trees promote wellbeing in diverse ways, from physical relief through heat mitigation to the mental health benefits of increased physical activity and reduced anxiety/depression (Remme et al., 2021). Benefits of urban trees are often determined by the composition of tree species, size, and density (Qiu et al., 2018; Villamagna et al., 2013; Ziter et al., 2013). The long-lived nature of trees means their current state, and the functioning of forest ecosystems, is strongly affected by legacy effects of past management decisions (Roman et al., 2021, 2018). Legacy effects may influence tree size and diversity due to legacies in soil properties and nutrient levels that continue to influence tree growth and functioning today (Aznarez et al., 2023; Pregitzer and Bradford, 2023). For example, agricultural land may be depleted of nutrients following its use as farmland, influencing the soil structure and development over time (Pabst et al., 2016). Indeed, agricultural land often accumulates carbon in the soil following development whereas former forest remnants do not, which has the potential to bring about long-term changes in tree growth and associated ecosystem services (Raciti et al., 2011). Legacy effects from past land-use types can also determine which species may or may not be present in an area (Munteanu et al., 2015). For example, in residential areas in Haikou, China that were previously woodlands, there are 7 species of trees that are not found in residential areas that were previously open land (Yang et al., 2017).

An urban park is thus a complex web of past and current land-uses and understanding the physical influences of historical land-use on vegetation will allow us to tease out a piece of the puzzle when assessing the differences in our parks’ benefits across space. Many cities are interested in pursuing restoration and altered management of urban green spaces. Quantifying the legacy effects of past land-use types on the benefits of trees in current urban parks will facilitate improved planning for equitable distribution of park benefits in the future. Here, we ask two research questions. First, what are the effects of historical land-use on current park forest structure, diversity, and consequently the capacity to provide cooling? Second, how do surrounding communities differ around parks of each historical land-use type, and what are the implications for equitable access to cooling?

We hypothesize that the historical land-use type will influence the quantity of temperature mitigation currently delivered by urban trees in Montreal’s parks via differences in tree size, tree basal area, and tree species richness. Tree size and basal area are both factors that drive cooling in urban settings (Helletsgruber et al., 2020; Rahman et al., 2020). Additionally, tree species is a driving determinant in cooling efficacy (Helletsgruber et al., 2020) and higher species richness of trees can lead to an increase in cooling through an increase in canopy density (Gillerot et al., 2024). We predict that previously industrial sites will have the lowest levels of temperature mitigation due smaller trees, lower basal area, and less species rich tree stands caused by high levels of heavy metals and low levels of carbon and nitrogen stocks, caused by soil contamination, removal, and treatment (Ashwood et al., 2022; Clarke et al., 2015; Goyal et al., 2020; Rooney et al., 2012). We predict previously agricultural sites will have higher capacity for temperature mitigation than previously industrial sites but less than previously forested sites. Agricultural lands typically have higher levels of heavy metals and lower carbon stocks than forested sites, which can cause a decrease in woody plant species richness (Arao et al., 2010; McLauchlan, 2006; Shumi et al., 2018; Zwolak et al., 2019) However, they can have high levels of nitrogen stocks, which increases plant growth (McLauchlan, 2006). Finally, we predict that previously forested sites will have the largest, densest trees and highest species richness, leading to the highest capacity for temperature mitigation due to their relatively less intense management (see Fig 1 for hypothesized causal relationships).

**Figure 1.**
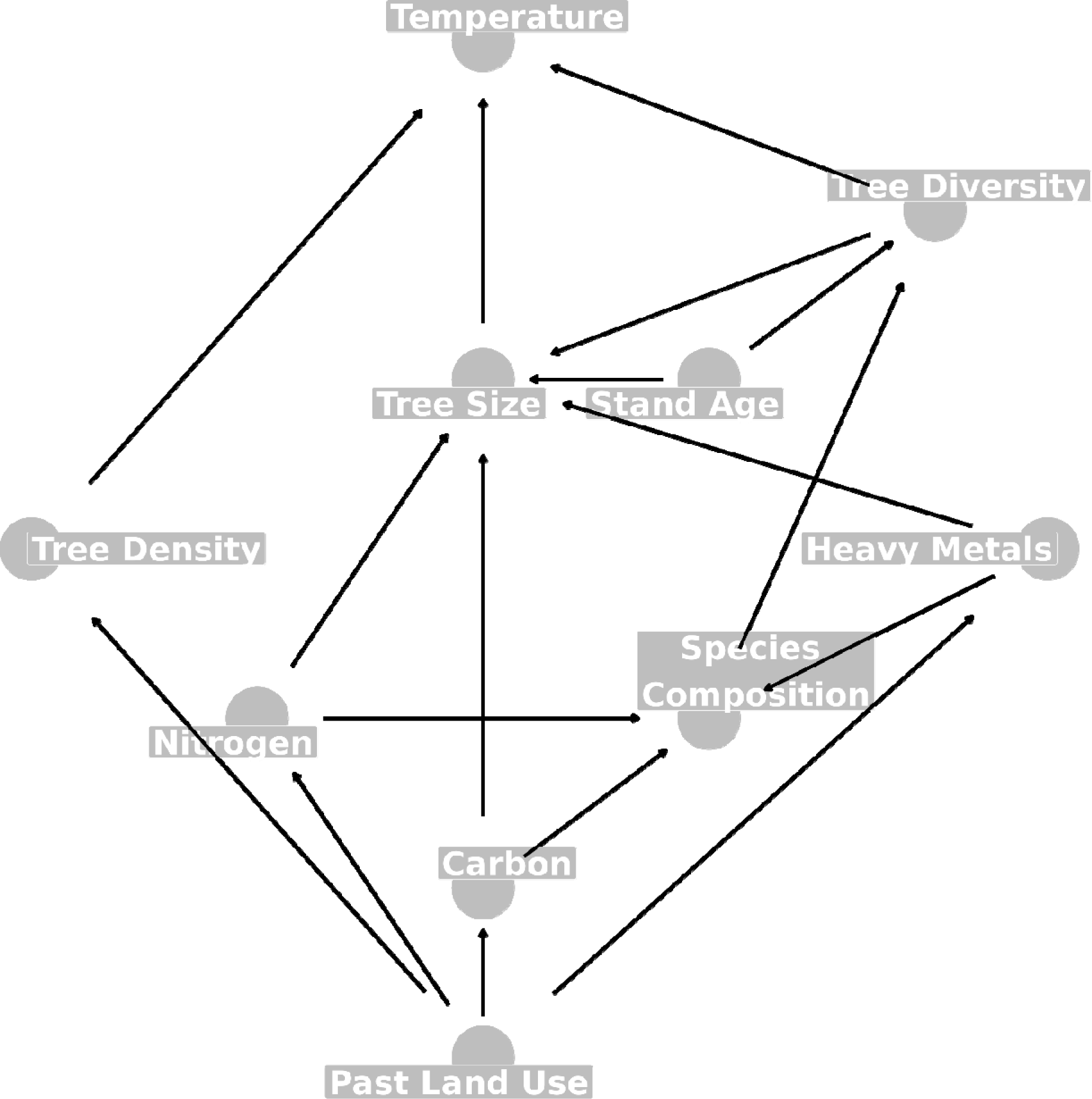
Directed acyclic graph depicting our hypothesized causal relationships between variables in our study system. Arrows in this graph indicate that the variable at the blunt end of the arrow directly affects the variable at the sharp end of the arrow (i.e., a causal relationship). For example, changing the past land-use will result in a change in nitrogen stocks. Temperature, tree density, tree size, tree diversity, past land use, and park age were all included in our models.

Regarding q uestion two, we hypothesize that due in part to historical environmental injustice there will be differences in resident communities surrounding parks characterised by each past land-use type. We predict that proportions of marginalized populations, such as recent immigrants, people of colour, and residents with low incomes will be present at higher rates surrounding parks that have industrial land-use legacies (Gelobter, 1993; Gerrish and Watkins, 2018; Jolivet and Carré, 2017; Landry et al., 2020). Due to the complex nature of these relationships, as well as historical data limitations, we do not have the ability to test the causal relationships between past land-use and current sociodemographic profile. However, we assess current differences among areas with different past land-use types using an exploratory data analysis so that we can better contextualize our results in terms of the capacity of cities to work towards green equity.

## 2 Methods

### 2.1 Study Area

We sampled 33 sites spread across the island of Montreal (hereafter referred to as “Montreal”) in the summers (May – August) of 2022 and 2023. The island of Montreal houses Canada’s second largest city, has 2.04 million residents (Gouvernement du Québec, 2022) and is approximately 4,022 km^2^. The island is composed of the 19 boroughs of the city of Montreal, along with 14 independent municipalities (Association des municipalités de banlieue, 2018; Ville de Montréal, 2023a), and the level of urbanization, population density, socioeconomic status, and development varies greatly within and across these areas. The parks that we sampled spanned the island (Fig. 2), including 13 boroughs and three municipalities.

**Figure 2.**
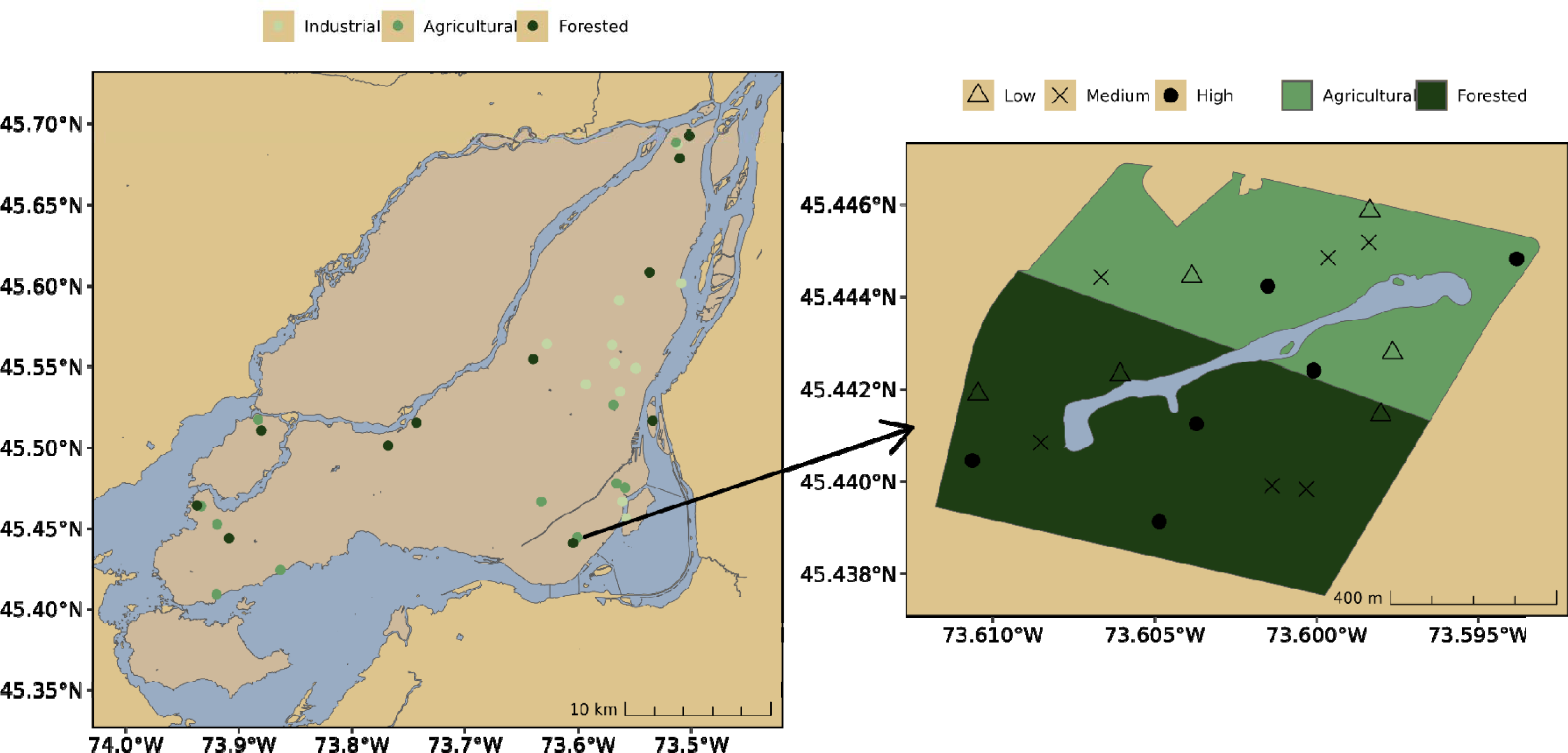
Lef t: Our study sites across the island of Montreal (n = 33, n = 11 per past land-use type). Different past land-uses were categorized into three main types, shown in different colour shades: industrial, agricultural, or forested. Right: Park Angrignon, one of our parks that had multiple previous land-use types. Each previous land-use type was considered a different study site and was sampled separately. Each study site had 15 potential sampling points (9 true sampling points), five in low canopy areas, five in medium canopy areas, and five in high canopy areas. For visual depiction of the sampling plots, see Supplementary Figure S1. Maps were created using ‘sf’, ‘ggplot2’, ‘MetBrewer’, and ‘osmdata’ packages in R 4.3.1 (Mills, 2022; Padgham et al., 2023; Pebesma et al., 2023a; R Core Team, 2023; Wickham et al., 2023)

### 2.2 Explanatory Variables

#### 2.2.1 Legacy Effects

To test how legacy effects influence park ecosystem service capacity, we identified the historical land-use type of 27 parks. We define historical land-use type as the category of land-use that the park was developed on and is therefore dependent on the year the park was developed, which ranges in our study from 1887 to 2016 (Table S2). Through extensive searching of historical city archives, satellite imagery, and grey literature published by historical societies, municipalities, and historical enthusiasts, we reconstructed the history of each park to identify a) the age of the park, and b) the land-use type immediately prior to park development (Table S1). We sorted historical land-uses into three categories: agricultural, industrial, and forested. We sampled 27 parks, however, within parks each different historical land-use was treated as a different study site, resulting in 33 study sites total (n = 11 of agricultural, industrial, and forested historical land-uses each, see Table S2 for historical land-use(s) of all parks). For parks with multiple historical land-uses, we digitized the different historical land-use boundaries using satellite imagery (Division de la géomatique and Ville de Montréal, 2023) and historical map layers from the City of Montreal (Division Soutien Technique, Infrastructures, CESM and Ville de Montréal, 2023; Ville de Montréal, 2023b) using QGIS v3.16 (QGIS Development Team, 2020)

#### 2.2.2 Forest Structure

We assessed forest structure of our study sites in the summer of 2022 by assessing tree basal area, median size, and species richness. To sample each site representatively, we sampled plots in areas of low, medium, and high canopy. Prior to entering the field, we calculated the canopy cover across the parks in 0.04 ha pixels (Communauté métropolitaine de Montreal, 2019). We then selected three low (10-30%), medium (31-80%), and high (81-100%) plots in each site to sample. If there were not enough low, medium, or high canopy areas to sample, we did as many as possible (n = 9 for study sites, with some exceptions when not possible, n = 279 sampling plots total). For each of the sampling plots, we conducted a forest inventory. Once arriving at the sampling plot, we walked 20 m in each cardinal direction and flagged the edge of the plot. The four cardinal points were then connected diagonally, creating a 0.08 ha square plot (Supplementary Figure S1). We measured and identified every tree with a diameter at breast height (DBH) > 5.0 cm. For the first half of the field season, we also did a complete inventory of the 0.08 ha plot for trees with a DBH > 1.0 cm. Due to time constraints, we changed our methodology to a “mini-plot” of 0.005 ha for trees between 1-5 cm DBH halfway through the season, consistent with the Canadian Forest Inventory (Canadian Forestry Inventory Committee, 2008). The mini-plot was placed in the most representative area of the larger sampling plot, and every tree under 5 cm DBH was identified and classified into a size category (1-3 cm DBH or 3-5 cm DBH, for more information see Figure S1). For trees with multiple stems, if the split in stems was above 30 cm we measured the DBH just below the split. If the split was below 30 cm, we measured each stem individually at breast height (Magarik et al., 2020). To identify trees to species level, we used National Audubon Society Field Guide to Trees: Eastern Region (Little, 1980), Trees of Canada (Farrar, 1995), and Seek (iNaturalist, 2022). If identifying to species level was not possible, we identified to the genus level (< 0.1 % of trees).

### 2.3 Response Variables

#### 2.3.1 Temperature Regulation

We collected temperature data the summer following the vegetative data collection. We deployed sensors during the hottest months of the year, collecting data between June 1 and July 31, 2023, recording a data point every 15 minutes. Sensors were CredoSense CSL-T0.5 temperature loggers, with an accuracy range of +/-0.1-0.5°C (Supplementary Figure S2). We placed temperature sensors in three plots within each site – one for each canopy level (Table S2). We installed sensors at approximately 1.5 m in height, in a location representative of the canopy cover of the plot, typically near the plot center, attached to a tree branch or trunk if the tree was too large to reach the branches.

To assess the cooling effect of park vegetation as an ecosystem service, instead of raw values of temperature, we deployed control sensors in the neighbourhoods surrounding each study site and compared the temperature *differences* between the park and the surrounding built-up area. We selected control sensors to be at least 500 m away from study sites or other large greenspaces (Hamada and Ohta, 2010; Vaz Monteiro et al., 2016). Control sensors were attached to poles in parking lots. If two study sites were very close together, and therefore have the same surrounding neighbourhood, only one control sensor was deployed for both sites. In total, we deployed 97 park sensors and 18 control sensors, for a total of 115 sensors. After collection, three sensors were malfunctioning and 14 were no longer at the site when we went to collect them, resulting in 98 full sensor datasets (see Supplementary Table S2 for sensor location details). One agricultural site had all three sensors stolen, resulting in 11 sites with temperature data for previously forested and previously industrial land-use legacies, and 10 sites for previously agricultural. To deal with missing control sensors, we supplemented our data using a dataset collected in the same summer, using the same type of sensor, with similar criteria to us or used the data from the next closest control site. For the dataset provided to us by a similar project, the control sensor was placed on a roadside, not in a parking lot. We conducted a sensitivity analysis to ensure that the data followed the same trends as other control sensors in our study (Figure S3).

#### 2.3.2 Neighbourhood Demographics

We calculated the neighbourhood demographics surrounding each study site using census data. Based on previous work on park access in Quebec (Apparicio et al., 2010; Robillard et al., 2023), we selected 1000 m as our buffer distance to include residents who may use the park regularly. Using census variables, we calculated population density, median income, proportion of people who are low income, recent immigrants, visible minorities, and those who have a bachelor’s degree or higher for the neighbourhoods within 1000 m surrounding each of our study parks at the dissemination area level (Government of Canada, 2021; Pham et al., 2013; Statistics Canada, 2022). When dissemination areas crossed buffer boundaries, we calculated the area that intersected the buffer and used a spatially weighted mean for calculations. For each past land-use type, we calculated the mean of each of our variables of interest within the industrial, agricultural, and forested groups. We then collected the data for each of our variables of interest at the broader island level (Statistics Canada, 2022). To obtain the difference between communities surrounding parks of each previous land-use type and the island of Montreal averages, we subtracted the values for the island from the mean values for the communities surrounding previously industrial, agricultural, and forested areas.

### 2.4 Statistical Analysis

We calculated tree species richness, basal area, and size composition at the plot level. We calculated all forest structure indices separately for trees < 5.0 cm DBH and trees > 5.0 cm DBH. Basal area was calculated by summing the cross-sectional area of each tree at breast height and then dividing the total by the plot size for a final unit of m^2^/ha. By calculating basal area instead of stem abundance, we were able to compare the density of trees < 5.0 cm even when plot size differed as they shared a common unit (m^2^/ha). Size composition was calculated by assessing the median DBH across all trees within a plot. For mini-plots, where trees < 5.0 cm DBH were classified into bins of 1-3 cm and 3-5 cm, for each recorded tree we randomly sampled values within the size categories to produce a dataset of real numbers. We only calculated species richness for trees >= 5 cm DBH due to the differences in sampling methodologies across the season for trees < 5 cm DBH. Since all plots were sampled at 100% coverage, we calculated species richness by summing the number of unique species found at each site.

To assess the cooling benefits provided by each plot, we calculated average temperature of daytime and nighttime hours per date for each sensor. Using the date, latitude, and longitude of each sensor, sunrise and sunset times were calculated via the ‘suncalc’ package (Thieurmel and Elmarhraoui, 2022). Sensors logged temperature every 15 minutes and we classified each logged point as daytime, if it was between sunrise and sunset, or nighttime, if it was before sunrise and after sunset. We then grouped the data by plot and date and calculated a mean temperature for daytime and nighttime. We did the same for each of the control sensors. We then calculated the cooling effect, separated by day and night for each date of sampling (n = 64 dates total). The cooling effect was calculated by taking the average temperature of each plot per day and subtracting the average temperature of the park’s control site for that day. We built our models using the causal framework outlined in Fig. 1. We used 5 Bayesian general linear mixed effects models to assess the effects that we were interested in (full model equations found in Supplementary Figure S4). We opted for Bayesian models due to their flexibility, generative nature, and production of probability distributions for effect sizes and comparisons instead of point estimates. To assess the total effect of past land-use on cooling effect, we modelled cooling effect as the response, with past land-use type time of day, and age of park as fixed effects. We also had an interaction effect between time of day, i.e., nighttime or daytime and past land-use type. Plot and park identification code (ID) were varying effects. To assess the total effects of past land-use type on forest structure, we produced three models with tree basal area, tree size, and tree diversity as response variables and past land-use type as a fixed explanatory variable for each. Then, to test the direct effects of forest structure on cooling, we modelled cooling effect as the response, with tree basal area, tree size, and tree diversity as fixed effects. We also included time of day, i.e., nighttime or daytime, as an interacting effect with each of the tree variables. Plot and park ID were included as varying effects. For all models, numeric variables were centered and scaled before modelling and weakly informative priors were selected. We define weakly informative priors as priors that increase the probability of biologically possible values for our variables while still allowing the possibility of extremely strong relationships. Setting weakly informative priors can improve chain convergence without affecting model performance or interpretation (McElreath, 2020). For our models with tree structure as response variables, we computed the global mean contrasts between each group (i.e., agricultural, forested, industrial) to assess differences between past land-uses using the posterior distributions of each model. For temperature models, the continuous explanatory variables in the model were set to their average values when computing the contrasts. We performed prior predictive checks using ‘tidybayes’ (Kay and Mastny, 2023) and ‘ggplot2’ (Wickham et al., 2023) packages to ensure that priors were appropriate (Supplementary Figure S5). We assessed model convergence by ensuring that all Rhat values were below 1.01, verifying that Markov chain Mont Carlo (MCMC) chains appeared well mixed through inspection of trace plots, and confirmed that there were minimal divergent transitions (< 0.5 %). To check if models were well specified, posterior predictive checks were completed. All model diagnostics can be found in Supplementary Figure S6. Data preparation and cleaning, spatial analysis, and modelling was done using R 4.3.0 (Bürkner et al., 2023; Pebesma et al., 2023a, 2023b; R Core Team, 2023).

## 3 Results

### 3.1 Overview

We sampled temperature across 61 days from June 1 to July 31, 2023. The mean temperature difference between park and control sites averaged across all days was 1.36 ± 1.0, with park plots nearly always cooler than control sites (98% of measurements). The cooling effect per day across all plots ranged from -0.8 to + 7.8, with a positive cooling effect indicating park plots providing more cooling than control sites. For forest structure variables we measured a total of 11,053 trees across all parks. For trees >= 5 cm diameter at breast height (DBH) at the plot scale, mean basal area ranged from 0.29 to 55.0 m^2^/ha, median size ranged from 6.4 cm DBH to 64.6 cm DBH, and mean species richness ranged from 1 to 11 species per park. Trees < 5 cm DBH ranged in basal area from 0.0 to 7.9m^2^/ha and median tree size ranged from 1.1 to 4.6 cm DBH.

### 3.2 Total Effect of Past Land-Use on Cooling

We found some differences in cooling effect across land use legacies (Figure 3). All past land-use types cooled more in during the day than at night (Figure 3). Previously forested plots cooled the most overall, with a mean daytime cooling effect of 1.5 ± 0.08 and a mean nighttime cooling effect of 1.4 ± 0.08. Previously agricultural sites were similar, with a mean daytime cooling effect of 1.4 ± 0.08 and a mean nighttime cooling effect of 1.3 ± 0.08. Previously industrial sites had the largest difference between daytime and nighttime cooling effects. Their daytime cooling effects were comparable to previously forested and agricultural sites, with a mean of 1.4 ± 0.09. However, previously industrial sites had the lowest nighttime cooling effects with a mean of 1.0 ± 0.09 (Figure 3).

**Figure 3.**
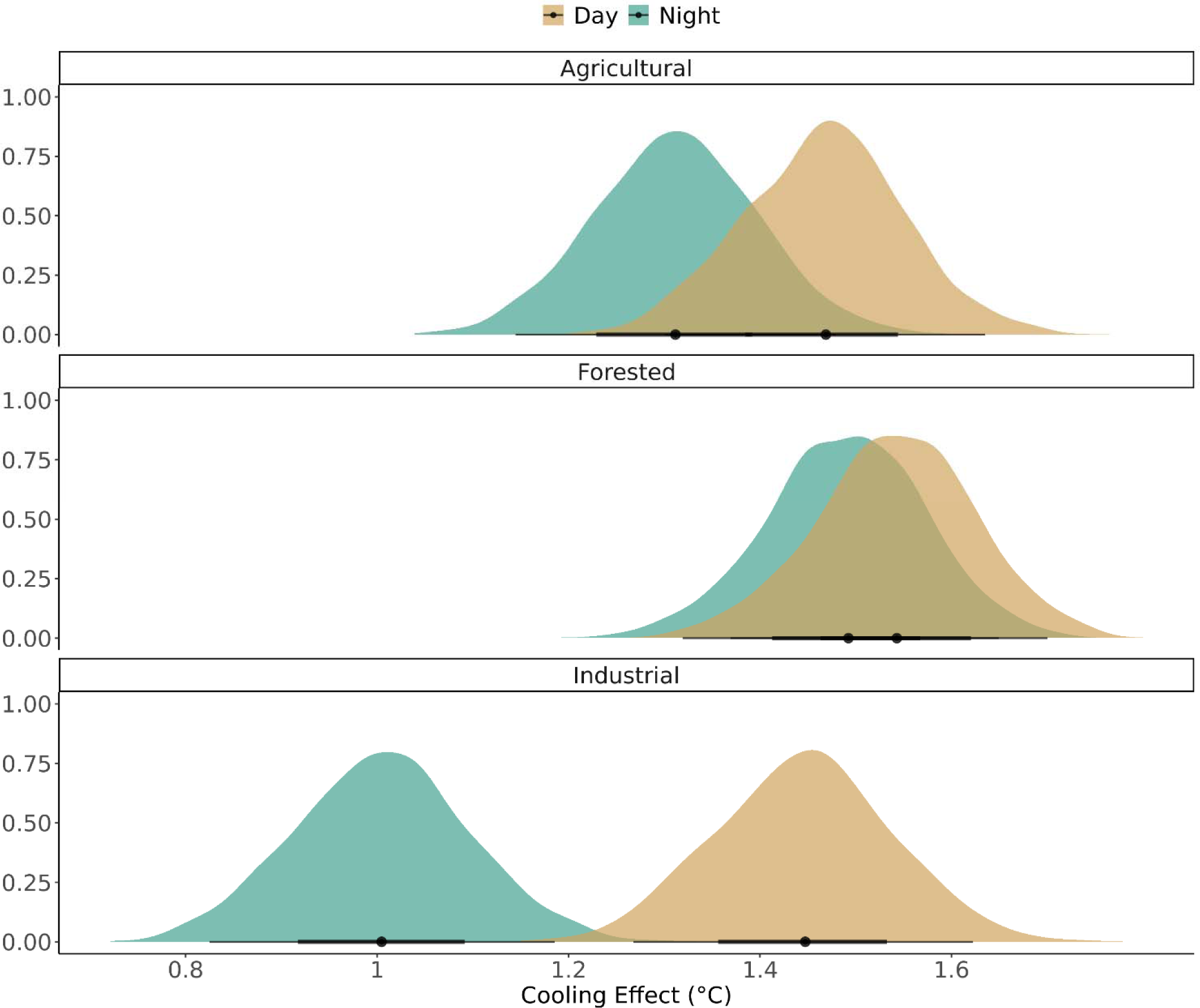
Visualization of the posterior draws for the model assessing total effect of past land-use on cooling effect of parks. Y-axis indicates density of the probability distributions, x-axis indicates relative cooling effect which is the cooling effect relative to the average cooling effect across the entire dataset Dot represents mean, linear intervals indicate 66% and 95% credible intervals. Night and daytime averages were calculated by categorizing each temperature measurement into day or night categories based on sunrise and sunset times of that day. Temperature was collected over 61 days in 2023.

### 3.3 Direct Effect of Past Land-Use on Forest Structure

We observed some differences in forest structure across different past land-use types (Figure 4). For large trees (DBH >= 5 cm), previously forested sites had the highest tree basal area. Previously forested sites had basal areas that were on average 0.72 ± 2.46 m^2^/ha higher than previously agricultural sites and 2.72 ± 2.80 m^2^/ha higher than previously industrial sites. Previously industrial sites had the largest trees >= 5 cm DBH, with a median DBH 8.8 ± 3.6 cm higher on average than previously forested sites and 3.9 ± 3.4 cm higher on average than previously agricultural sites. Previously agricultural sites were 4.8 ± 3.4 cm larger on average than previously forested sites (Figure 4). Similar to large tree basal area, species richness was highest in previously forested sites but differences were small overall (Figure 4) Previously forested sites had on average 0.8 ± 0.5 more species than previously agricultural sites and 1.1 ± 0.5 more species than industrial sites. Previously agricultural sites had on average 0.3 ± 0.5 more species than previously industrial sites. Regarding small (DBH < 5 cm) basal area, previously agricultural sites had the highest basal area, with on average 0.56 ± 0.49 m^2^/ha higher basal area than previously forested sites and 0.22 ± 0.55 m^2^/ha higher basal area than previously industrial sites. Previously industrial sites had higher small tree basal area than previously forested sites, on average 0.33 ± 0.67 m^2^/ha higher. Consistent with large trees, we found that previously industrial sites had the largest small trees (Figure 4). Previously industrial sites had on average 0.5 ± 0.2 cm larger trees than previously forested sites and 0.2 ± 0.2 cm larger trees than previously agricultural sites. Previously agricultural sites had 0.3 ± 0.2 cm larger trees than previously forested sites on average.

**Figure 4.**
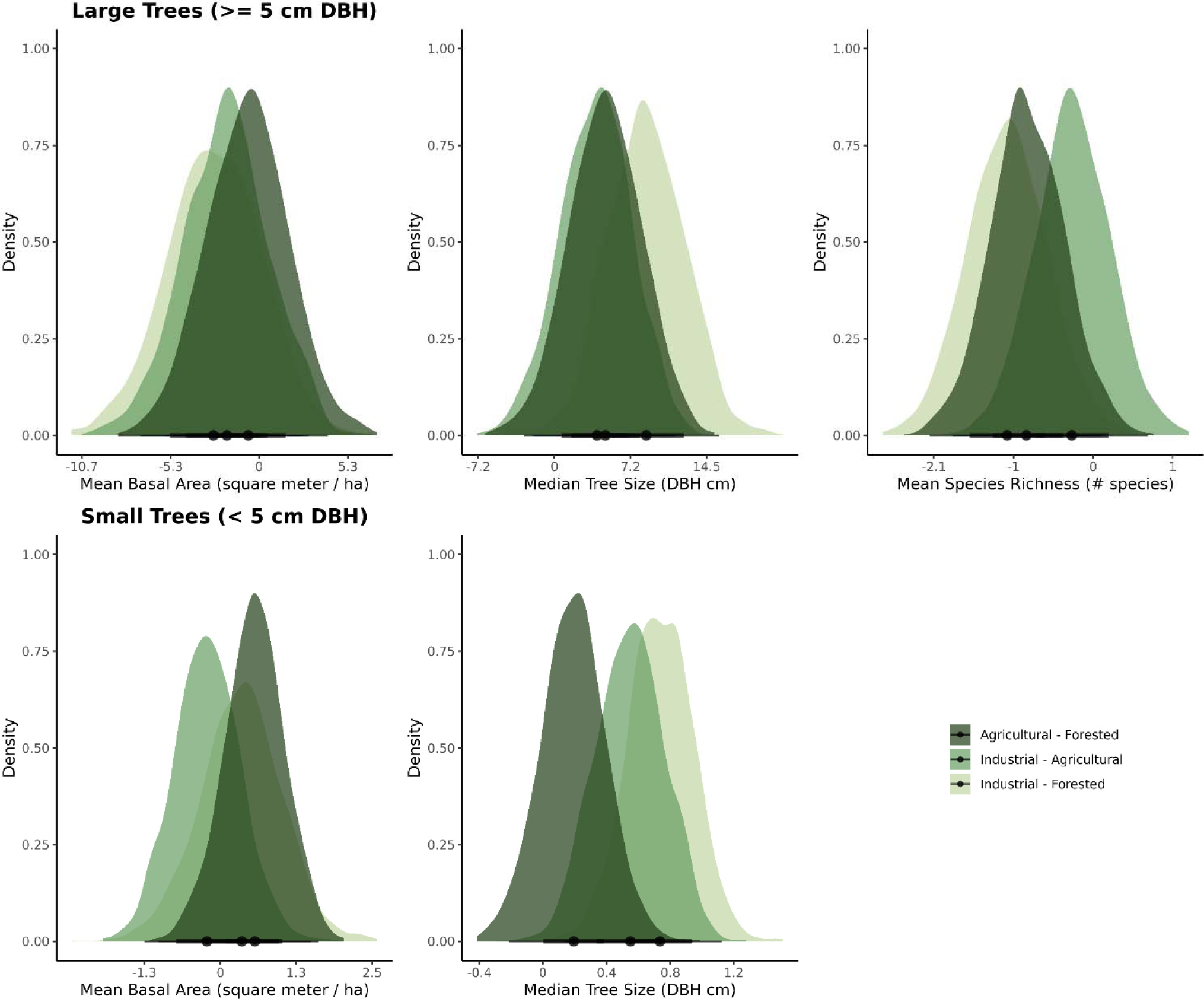
Visualization of the pairwise contrasts of the posterior draws for each combination of past land-use type to assess the direct effects of past land-use on forest structure of parks. Top 3 panels show the relationships between cooling and forest structure of large trees (trees with a diameter at breast height (DBH) of >= 5 cm). Bottom two panels show the relationships between cooling and forest structure of small trees (trees with DBH < 5 cm). Dot represents global mean of the contrast, linear intervals indicate 66% and 95% credible intervals. Contrasts can be interpreted as the effect of switching from the state on the right, e.g., forested, to the state on the left, e.g., agricultural.

### 3.4 Direct Effect of Forest Structure on Cooling

We found evidence that differences in forest structure alter the levels of cooling provided by parks (Figure 5). For trees >= 5 cm DBH, we found a positive effect between tree basal area and cooling during the day and a negative effect between tree basal area and cooling at night. Large tree size and species richness had little to no effect on cooling. For trees < 5 cm DBH, the relationship between basal area and cooling switches – we found a positive relationship between small tree basal area and cooling during nighttime and a slightly negative effect between small tree basal area and cooling during the day. Small tree size had a weak positive relationship with cooling during the day and a weak negative relationship with cooling during the night (Figure 5).

**Figure 5.**
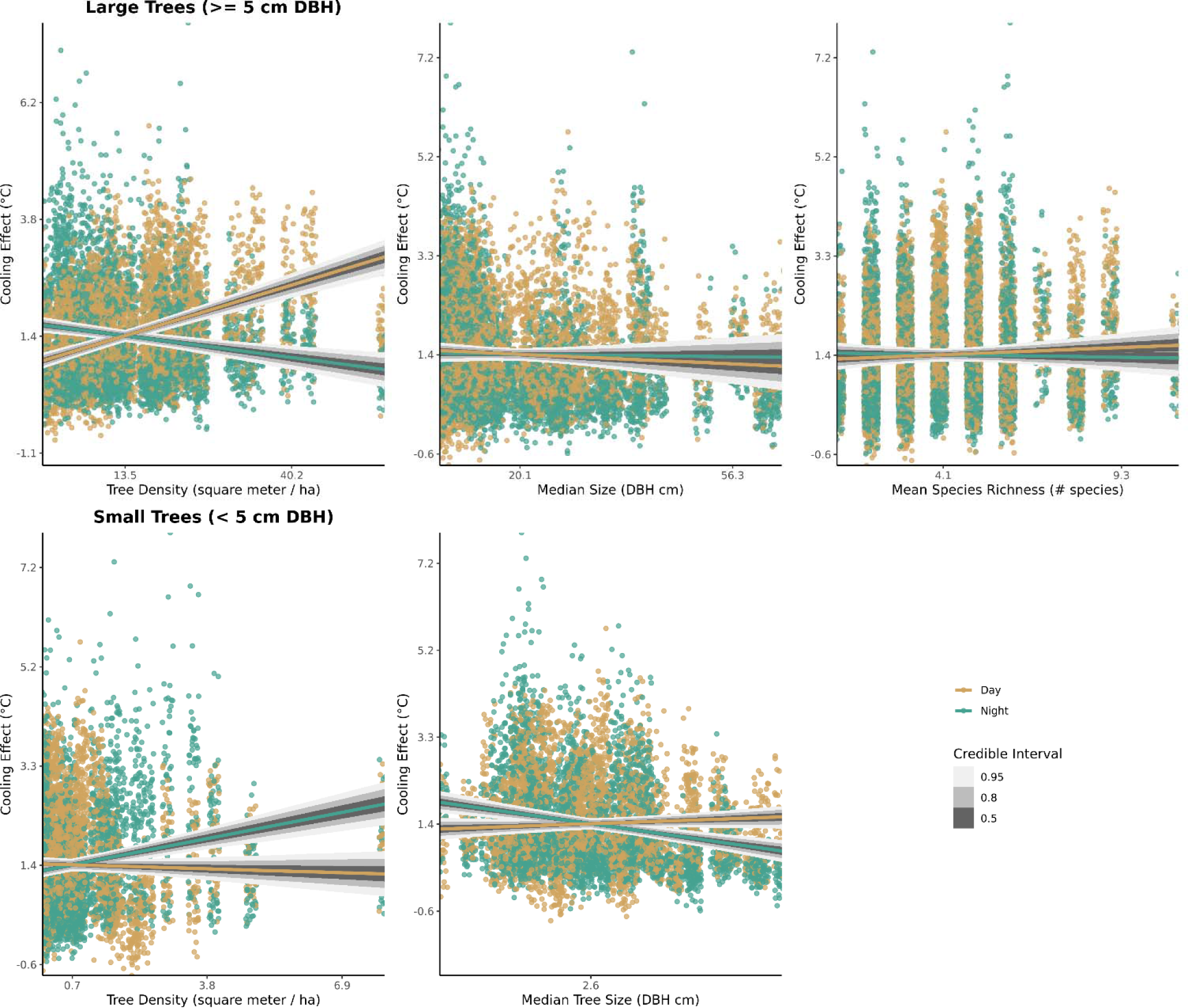
Visualization of the predicted relationships between forest structure and relative cooling effect, calculated while holding all other variables in the model constant and using time of day as an interaction effect. Predicted draws were calculated 1,000 times, and then extracted and plotted. Coloured lines indicate the conditional expectation of the model to the data, with grey credible intervals at 0.50, 0.8, and 0.95. Top 3 panels show the relationships between cooling and forest structure of large trees (trees with a diameter at breast height (DBH) of >= 5 cm). Bottom two panels show the relationships between cooling and forest structure of small trees (trees with DBH < 5 cm). Points represent raw data, coloured by time of day.

### 3.2 Neighbourhood Sociodemographic Composition

Due to data limitations, we were only able to offer a qualitative description of the communities surrounding our sites using current census data. Using these data, we did find differences among communities surrounding past land-use types. However, we also found evidence for general systemic inequity in the immediate areas surrounding our parks, consistent with previous literature (Table 1). There was a higher proportion of residents with bachelor’s degrees or higher, and a lower proportion of visible minorities in the areas immediately surrounding parks of all historical legacies than we would expect given the island of Montreal’s averages (Table 1). Using census data, we found trends that support what we would expect regarding legacy effects of environmental injustice when looking at the areas surrounding previously industrial parks. Areas surrounding previously industrial parks had a lower median income and prevalence of single detached homes, and the highest population density when compared to Montreal’s averages and other land-use types.

**Table 1.** Mean difference in socioeconomic variables between neighbourhoods surrounding each park type and island of Montreal’s averages, for parks with each of three different historical land-uses. Positive values indicate a value higher than the island’s average and negative indicate a value lower than the island’s average. Standard deviations are presented beside means.

| Past Land Use | Population Density (per km2) | Median Total Income (\$) | Single Detached Homes (% residents) | Visible Minorities (% residents) | Recent Immigrants (% residents) | Attained a Bachelor's Degree or Higher (% residents) |
| --- | --- | --- | --- | --- | --- | --- |
| Industrial | 7683 +/- 2576 | -1849 +/- 2440 | -7.3 +/- 7.5 | -10.4 +/- 10.4 | 3.5 +/- 2.0 | 6.8 +/- 13.6 |
| Agricultural | 2537 +/- 6041 | 1230 +/- 3855 | 28.3 +/- 39.6 | -8.6 +/- 9.9 | 2.1 +/- 2.9 | 14.4 +/- 11.0 |
| Forested | 1571 +/- 3498 | 698 +/- 2908 | 21.2 +/- 28.6 | -3.4 +/- 11.0 | 3 +/- 3.1 | 5.1 +/- 12.2 |

## 4 Discussion

We leveraged historical archives, urban forestry fieldwork, and sensor data to ask two research questions in Montreal, QC. For our first question, we found support for our hypothesis that capacity for cooling differs across previous land-use types, most notably the cooling effect of previously industrial sites at nighttime is lower than at previously agricultural and previously forested sites. Further, we found that forest structure had some differences across land-use type. Previously forested parks had smaller trees with a higher basal area and a slight increase (∼ 1 species on average) in species richness. In contrast, previously industrial sites had lower basal area but larger trees on average and previously agricultural sites had the highest density of trees < 5 cm DBH.

### 4.1 Effect of Past Land-Use on Cooling

All park types show capacity for cooling, with similar, stronger levels of cooling capacity across park type during the daytime. A stronger daytime cooling effect is expected, as canopy cooling effects are typically stronger during the day (Wujeska-Klause and Pfautsch, 2020; Ziter et al., 2019). The similar levels of daytime cooling across past land-use type is a testament to park management and provides evidence that park development can provide a critical ecosystem service despite varied land-use histories. However, we observed a difference in cooling capacity across past land-use types at night. We hypothesize that the observed nighttime difference between parks with varying land-use legacies is reflecting the differences in the broader urban heat island effect, which is often felt most intensely at night (Iungman et al., 2023). Given that previously industrial sites are generally smaller parks located in denser urban areas, the reduced contributions of trees to cooling at night via lower transpiration rates and lack of shade provision (Ziter et al., 2019) mean that the parks in these areas are less able to combat the stronger urban heat island effect. This is further evidenced by the nighttime cooling effect observed in previously forested parks. Previously forested parks had dense tree stands >= 5 cm DBH which causes a reduction in nighttime cooling capacity (Figure 5), yet they still have the highest levels of nighttime cooling. Due to their large size and positioning in lower density areas, previously forested parks most likely experience less of the urban heat island effect and are therefore cooler at night despite their high tree density. Thus, we observe that land use legacy effects on cooling capacity may manifest not only via changes to forest structure, but also the landscape context in which we typically find parks with different historical land uses.

These nighttime differences in cooling capacity are especially important to understand as nighttime temperature is often considered a better predictor of mortality and morbidity (Sarofim et al., 2016). Our study demonstrated differences in cooling capacity, however, there are other microclimatic variables that impact people’s perception of heat and resulting heat-related illnesses, including relative humidity and wind, that were not included in this study. Further investigating how land-use legacies influence multiple microclimatic variables which affect the urban heat island effect and perceived heat would be a useful addition to the literature

### 4.2 Effect of Past Land-Use on Forest Structure

Forest structure also differed across past land-use types, especially in relation to basal area and tree size. Previously forested sites had smaller trees with higher basal area, which is consistent with what we would expect from the literature, as temporal distance from anthropogenic disturbance is positively correlated with basal area in other studies (Pregitzer and Bradford, 2023; Thompson et al., 2002). One limitation when working with historical data is often it is not possible to find the complete history of a site. As such, we were unable to determine the logging and management history of the previously forested sites, though the small, dense stands suggest that these areas are relatively young forests (Thompson et al., 2002). Previously industrial sites had the opposite trend to forested, characterized by lower basal area but higher median tree size when compared to other past land-use types. Previously industrial sites having larger trees may be due to the inherent nature of their establishment. Industrial sites have the topsoil replaced upon restoration and conversion to a park, and often establish their woodlands starting with intentional planting of saplings due to low natural recruitment (Ashwood et al., 2022; Robinson et al., 1992). Newly planted saplings in Montreal are generally required to be between 5-7 cm DBH, which are larger than many of the naturally establishing saplings in the previously agricultural and forested sites (Anjou, 2019; Rivière-des-Prairies–Pointe-aux-Trembles, 2013; Saint-Laurent, 2018). These results suggest that there is a density-size trade-off present in urban parks that is driving differences across past land-use types. This trade-off has been previously documented, as large trees can be difficult to maintain and manage especially in high volumes (Endreny et al., 2017; McPherson et al., 1997; Soto et al., 2018). Therefore, urban forests stands often alternate between two options, smaller, dense trees or larger, sparser trees as a strategy to balance size, density, and cover.

One similarity in forest structure that we found across past land-use types was species richness, where previously forested sites had slightly higher levels than previously industrial and agricultural sites but only by an average of ∼ 1 species. This is contrary to what we would expect from the literature (Aznarez et al., 2023; Pregitzer and Bradford, 2023; Roman et al., 2021, 2018). For example, species composition of New York forests is dependent on land-use history (Pregitzer and Bradford, 2023). However, we did not find any evidence of differences in species richness across sites. This finding is a testament to the power of park management, and shows that maintaining tree diversity is possible across a city with proper and cohesive management (Aznarez et al., 2022). However, we recognize that species richness is not always the most meaningful measure of diversity and indeed has no effect on cooling in our study. Future studies looking at how legacy effects influence functional diversity and the presence of native vs invasive species (Pregitzer and Bradford, 2023) would be an interesting extension of this research.

### 4.3 Effect of Forest Structure on Cooling

For trees >= 5 cm DBH, cooling is driven mainly by basal area of trees. We observe a positive relationship between cooling and basal area during the day, and a negative relationship between cooling and basal area during the night. This is what we would expect from the literature, higher tree density typically results in more tree cover and thus higher rates of transpiration and shade provision, which increases cooling during the day (He et al., 2021; Zhao et al., 2023; Ziter et al., 2019). At night, high tree cover can actually trap heat, decreasing the cooling effect (Wujeska-Klause and Pfautsch, 2020). However, we would expect this relationship to hold with tree size, and it does not. We see no relationship between tree size for trees >= 5 cm DBH and cooling. This is unexpected and contrary to our predictions, as there are many studies that demonstrate that larger size of trees in parks results in greater cooling (Helletsgruber et al., 2020; Rahman et al., 2020). We hypothesize that our contrary results are due to the density-size trade-off. Due to planting and maintenance strategies in urban park management, development patterns, and resource constraints, it is rare to find a park that has both large and dense trees (e.g., see Supplementary Figure S8). Further, the largest trees in urban parks are often surrounded by mowed turfgrass, leading to a mitigation of the cooling effect, especially on hot days (Kraemer and Kabisch, 2022; Rahman et al., 2020). In contrast, more “natural” or forest-like conditions in our study parks tended to have denser stands of small trees, but with more structural contrast. Our findings suggest that large trees alone are not enough to provide cooling in urban parks, with high tree density and/or cover providing stronger daytime cooling. Although we were not able to measure understory structure in our study, literature suggests that it can also impact cooling (Fung and Jim, 2019; Stickley and Fraterrigo, 2021) and should be considered in the future as a potentially important component of urban park cooling capacity.

Increasing the basal area of trees in parks or creating more “naturalized” or rewilded areas may be good strategies to provide temperature regulation during the day. However, dense, forested parks may provide daytime cooling at the expense of nighttime cooling and the many other needs that parks serve (Gaston et al., 2013; Kremer et al., 2016). Parks with only dense forest stands lose the opportunity to provide open space for recreation, community gardens for food provisioning, wildflower beds for pollinators, etc. To further tease out the impact that forest structure can have, we suggest looking at the effects of forest structure within parks. Understanding both within and between group variation will help inform the role of management on the capacity for cooling, alongside strategies to balance this benefit with other desired ecosystem services.

When looking at trees < 5 cm DBH, our results are unexpected. We found that as basal area of small trees increases, the nighttime cooling effect increases, and the daytime cooling effect is limited. This directly conflicts with what we would expect based on the literature and our results for trees >= 5 cm DBH. We hypothesize that this relationship is due a trade-off between large and small trees in our system. Areas with a high basal area of trees < 5 cm DBH – i.e., dense, shrubby stands – are less likely to contain a large shade tree in the parks we studied, therefore heat can escape at night and results in a positive relationship between basal area and nighttime cooling. However, we do see the expected relationship with tree size. An increase in median tree size causes an increase in daytime cooling and a decrease in nighttime cooling. This is what we expected to see given the literature, where larger trees increase daytime cooling because of increased shade provision but trap heat at night (Wujeska-Klause and Pfautsch, 2020).

### 4.4 Communities Surrounding Past Land-Use Types

Our second hypothesis was partially supported. We found that communities surrounding previously industrial parks had a lower median income and higher population density than expected compared to previously agricultural and forested parks and to the Montreal area average, indicating persistent effects of environmental inequities. Industrial areas in Montreal have a history of systemic inequity, with neighbourhoods surrounding areas that were used for quarrying, garbage dumping, and/or other industrial activities being inhabited by working class or low-income people, who are often recently immigrated, and are majority Black or Brown (Kikano et al., 2023; Schell et al., 2020; Waldron, 2018). Our results provide evidence that park establishment in these areas can provide critical daytime cooling, though the critical service of nighttime cooling is lacking. With management that incorporates strategies to improve nighttime cooling and maintain daytime cooling, parks established on previously industrial land have the potential to counter a history of inequity in these neighbourhoods, as they are providing important natural benefits to people with lower median incomes living in denser populations.

A natural conclusion to draw from our results is that previously industrial areas should be priority sites for development of new urban green spaces, to increase nighttime cooling capacity and combat the urban heat island effect. However, we are wary of making that recommendation, as we found that communities surrounding parks in all three of our past land-use types have lower proportions of visible minorities and higher proportions of highly educated people than average for the island of Montreal. This finding supports the luxury effect – an established relationship across North America between wealth/privilege and plant diversity (Hope et al., 2003) that has been expanded to include vegetation metrics beyond diversity such as greenness (Mennis, 2006), vegetation cover (Clarke et al., 2013), and more (Schell et al., 2020). This indicates to us that efforts to provide urban nature, involving the establishment or restoration of urban green spaces, may fail in their mission and result in psychological and physical displacement of the communities that they are posed to help (Anguelovski, 2016; Sax et al., 2022). Green gentrification is an issue that we commonly see in Canada and around the world, and which can only be solved by stronger housing regulations and community-led policies (Goossens et al., 2020; Parish, 2020; Triguero-Mas et al., 2021). Further investigation into the connections between historical injustice, ecological capacity of greenspaces, and gentrification is warranted. We recommend that moving forward, urban greening is done in collaboration with housing and urban planning departments, and driven by community-led greening initiatives.

In conclusion, we found that incorporating historical context can be a useful way understand differences in contemporary ecological and societal conditions. Building on this study by incorporating other key ecosystem services such as air pollution mitigation, recreation provision, cultural importance, and habitat provisioning is integral to discovering how our histories influence our current day service provision, as well as informing how our current decision making will influence our future service provision. We see through this work that the restoration of industrial sites has the potential to provide critical ecosystem services to equity-seeking groups and suggest this as a promising strategy to provide urban nature as long as it is done intentionally and led by the community it is serving. Finally, our results support that building greenspaces on all different types of sites, from forest remnants to quarries, results in the capacity for daytime cooling and comparable tree diversity, critical benefits to our cities.

## Acknowledgements

We gratefully acknowledge Adela Bartova, Angelina Teichrib, and Gabrielle Gingras for their field support. Thank you to Andrew MacDonald for help with modelling. We would also like to thank the cities of Montreal, Beaconsfield, and Bai-d’Urfé for their logistical support. Thank you to reviewers from the Ziter Urban Landscape Ecology lab for their invaluable edits to earlier versions of this manuscript. Finally, thank you to several anonymous reviewers for their constructive feedback.

## Funding

This research was funded by the Natural Sciences and Engineering Research Council of Canada (CGSD3-559539-2021 to ICR and Discovery Grant RGPIN-2019-07010 to CDZ); UFOR NSERC CREATE scholarships awarded to EP and MAP; and a Mitacs Globalink Internship awarded to LCGS.

## Conflict of Interest

The authors declare that there is no conflict of interest.

## Data Availability Statement

All data and code for sampling, data cleaning, and analysis can be found at https://github.com/zule-lab/ParkLegacies, archived at https://zenodo.org/records/10406628.

## Notes

https://zenodo.org/records/10406628

### Competing Interest Statement

The authors have declared no competing interest.

## Literature Cited

Alexander, E.R., 1992. Approaches to Planning: Introducing Current Planning Theories, Concepts and Issues, 2nd ed. Gordon and Breach Publishers.

Alig, R.J., Kline, J.D., Lichtenstein, M., 2004. Urbanization on the US landscape: looking ahead in the 21st century. Landscape and Urban Planning 69, 219–234. 10.1016/j.landurbplan.2003.07.004

Anguelovski, I., 2016. From Toxic Sites to Parks as (Green) LULUs? New Challenges of Inequity, Privilege, Gentrification, and Exclusion for Urban Environmental Justice. Journal of Planning Literature 31, 23–36. 10.1177/0885412215610491

Anguelovski, I., Irazábal Zurita, C., Connolly, J.J.T., 2019. Grabbed Urban Landscapes: Socio-spatial Tensions in Green Infrastructure Planning in Medellín. International Journal of Urban and Regional Research 43, 133–156. 10.1111/1468-2427.12725

Anjou, 2019. Plan Maitre de plantation de l’arrondissement d’Anjou 2019-2023. Ville de Montréal.

Apparicio, P., Cloutier, M.-S., Séguin, A.-M., Ades, J., 2010. Accessibilité spatiale aux parcs urbains pour les enfants et injustice environnementale. Exploration du cas montréalais. Revue internationale de géomatique 20, 363–389. 10.3166/rig.20.363-389

Arao, T., Ishikawa, S., Murakami, M., Abe, K., Maejima, Y., Makino, T., 2010. Heavy metal contamination of agricultural soil and countermeasures in Japan. Paddy Water Environ 8, 247–257. 10.1007/s10333-010-0205-7

Ashwood, F., Barreto, C., Butt, K.R., Lampert, M., Doick, K., Vanguelova, E.I., 2022. Earthworms and soil mesofauna as early bioindicators for landfill restoration. Soil Res. 61, 311–328. 10.1071/SR21286

Association des municipalités de banlieue, 2018. Municipalities [WWW Document]. URL https://www.amb-asm.qc.ca/en/municipalities/ (accessed 10.17.23).

Aznarez, C., Svenning, J.-C., Pacheco, J.P., Have Kallesøe, F., Baró, F., Pascual, U., 2023. Luxury and legacy effects on urban biodiversity, vegetation cover and ecosystem services. npj Urban Sustain 3, 1–11. 10.1038/s42949-023-00128-7

Aznarez, C., Svenning, J.-C., Taveira, G., Baró, F., Pascual, U., 2022. Wildness and habitat quality drive spatial patterns of urban biodiversity. Landscape and Urban Planning 228, 104570. 10.1016/j.landurbplan.2022.104570

Bürkner, P.-C., Gabry, J., Weber, S., Johnson, A., Modrak, M., Badr, H.S., Weber, F., Ben-Shachar, M.S., Rabel, H., Mills, S.C., Wild, S., 2023. brms: Bayesian Regression Models using “Stan.”

Calmore, J.O., 1997. Race/ism lost and found: The fair housing act at thirty. University of Miami Law Reviews 52, 1067.

Canadian Forestry Inventory Committee, 2008. Canada’s National Forest Inventory Ground Sampling Guidelines.

Chao, A., Gotelli, N.J., Hsieh, T.C., Sander, E.L., Ma, K.H., Colwell, R.K., Ellison, A.M., 2014. Rarefaction and extrapolation with Hill numbers: a framework for sampling and estimation in species diversity studies. Ecological Monographs 84, 45–67. 10.1890/13-0133.1

Chao, A., Jost, L., 2012. Coverage-based rarefaction and extrapolation: standardizing samples by completeness rather than size. Ecology 93, 2533–2547. 10.1890/11-1952.1

Clarke, L.W., Jenerette, G.D., Bain, D.J., 2015. Urban legacies and soil management affect the concentration and speciation of trace metals in Los Angeles community garden soils. Environmental Pollution 197, 1–12. 10.1016/j.envpol.2014.11.015

Clarke, L.W., Jenerette, G.D., Davila, A., 2013. The luxury of vegetation and the legacy of tree biodiversity in Los Angeles, CA. Landscape and Urban Planning 116, 48–59. 10.1016/j.landurbplan.2013.04.006

Communauté métropolitaine de Montréal, 2019. Indice canopée métropolitain.

Cramer, M.D., Hawkins, H.-J., Verboom, G.A., 2009. The importance of nutritional regulation of plant water flux. Oecologia 161, 15–24. 10.1007/s00442-009-1364-3

Dallimer, M., Davies, Z.G., Diaz-Porras, D.F., Irvine, K.N., Maltby, L., Warren, P.H., Armsworth, P.R., Gaston, K.J., 2015. Historical influences on the current provision of multiple ecosystem services. Global Environmental Change 31, 307–317. 10.1016/j.gloenvcha.2015.01.015

Division de la géomatique, Ville de Montréal, 2023. Photothèque 1930-2003.

Division Soutien Technique, Infrastructures, CESM, Ville de Montréal, 2023. Carte de localisation des anciennes carrières et des dépôts de surface de la Ville de Montréal.

Endreny, T., Santagata, R., Perna, A., Stefano, C.D., Rallo, R.F., Ulgiati, S., 2017. Implementing and managing urban forests: A much needed conservation strategy to increase ecosystem services and urban wellbeing. Ecological Modelling 360, 328–335. 10.1016/j.ecolmodel.2017.07.016

Fan, J.Y., Sengupta, R., 2022. Montreal’s environmental justice problem with respect to the urban heat island phenomenon. Canadian Geographies /Géographies canadiennes 66, 307–321. 10.1111/cag.12690

Farrar, J.L., 1995. Trees in Canada. Fitzhenry & Whiteside.

Foster, D., Swanson, F., Aber, J., Burke, I., Brokaw, N., Tilman, D., Knapp, A., 2003. The Importance of Land-Use Legacies to Ecology and Conservation. BioScience 53, 77. 10.1641/0006-3568(2003)053[0077:TIOLUL]2.0.CO;2

Fung, C.K.W., Jim, C.Y., 2019. Microclimatic resilience of subtropical woodlands and urban-forest benefits. Urban Forestry & Urban Greening 42, 100–112. 10.1016/j.ufug.2019.05.014

Gaston, K.J., Avila-Jim Enez, L., Edmondson, J.L., 2013. Managing urban ecosystems for goods and services. Journal of Applied Ecology 50, 830–840. 10.1111/1365-2664.12087

Gelobter, M., 1993. The Meaning of Urban Environmental Justice. Fordham Urb. L.J. 21, 841–856.

Gerrish, E., Watkins, S.L., 2018. The relationship between urban forests and income: A meta-analysis. Landscape and Urban Planning 170, 293–308. 10.1016/j.landurbplan.2017.09.005

Gillerot, L., Rozario, K., de Frenne, P., Oh, R., Ponette, Q., Bonn, A., Chow, W., Godbold, D., Steinparzer, M., Haluza, D., Landuyt, D., Muys, B., Verheyen, K., 2024. Forests are chill: The interplay between thermal comfort and mental wellbeing. Landscape and Urban Planning 242, 104933. 10.1016/j.landurbplan.2023.104933

Goetz, E.G., Williams, R.A., Damiano, A., 2020. Whiteness and Urban Planning. Journal of the American Planning Association 86, 142–156. 10.1080/01944363.2019.1693907

Goossens, C., Oosterlynck, S., Bradt, L., 2020. Livable streets? Green gentrification and the displacement of longtime residents in Ghent, Belgium. Urban Geography 41, 550–572. 10.1080/02723638.2019.1686307

Gouvernement du Québec, 2022. Démographie [WWW Document]. Ministère de l’Économie, de l’Innovation et de l’Énergie. URL https://www.economie.gouv.qc.ca/pages-regionales/montreal/portrait-regional/demographie (accessed 10.17.23).

Government of Canada, S.C., 2021. 2021 Census Boundary files.

Government of Canada, Statistics Canada, 2023. Census of Population - Montréal, Territoire équivalent (TÉ) [Census division], Quebec.

Goyal, D., Yadav, A., Prasad, M., Singh, T.B., Shrivastav, P., Ali, A., Dantu, P.K., Mishra, S., 2020. Effect of Heavy Metals on Plant Growth: An Overview, in: Contaminants in Agriculture. Springer, Cham, pp. 79–101. 10.1007/978-3-030-41552-5_4

Grove, M., Ogden, L., Pickett, S., Boone, C., Buckley, G., Locke, D.H., Lord, C., Hall, B., 2018. The Legacy Effect: Understanding How Segregation and Environmental Injustice Unfold over Time in Baltimore. Annals of the American Association of Geographers 108, 524–537. 10.1080/24694452.2017.1365585

Haaland, C., van den Bosch, C.K., 2015. Challenges and strategies for urban green-space planning in cities undergoing densification: A review. Urban Forestry & Urban Greening 14, 760–771. 10.1016/j.ufug.2015.07.009

Hamada, S., Ohta, T., 2010. Seasonal variations in the cooling effect of urban green areas on surrounding urban areas. Urban Forestry & Urban Greening 9, 15–24. 10.1016/j.ufug.2009.10.002

He, C., Zhou, L., Yao, Y., Ma, W., Kinney, P.L., 2021. Cooling effect of urban trees and its spatiotemporal characteristics: A comparative study. Building and Environment 204, 108103. 10.1016/j.buildenv.2021.108103

Healy, M., Rogan, J., Roman, L.A., Nix, S., Martin, D.G., Geron, N., 2022. Historical Urban Tree Canopy Cover Change in Two Post-Industrial Cities. Environmental Management 70, 16–34. 10.1007/s00267-022-01614-x

Helletsgruber, C., Gillner, S., Gulyás, Á., Junker, R.R., Tanács, E., Hof, A., 2020. Identifying Tree Traits for Cooling Urban Heat Islands—A Cross-City Empirical Analysis. Forests 11, 1064. 10.3390/f11101064

Héroux, M., Martin, D., 2020. FREDERIC-BACK PARK, MONTREAL, CANADA: HOW 40 MILLION TONNES OF SOLID WASTE SUPPORT A PUBLIC PARK. Detritus 68–80. 10.31025/2611-4135/2020.13972

Hope, D., Gries, C., Zhu, W., Fagan, W.F., Redman, C.L., Grimm, N.B., Nelson, A.L., Martin, C., Kinzig, A., 2003. Socioeconomics drive urban plant diversity. Proceedings of the National Academy of Sciences 100, 8788–8792. 10.1073/pnas.1537557100

iNaturalist, 2022. Seek.

Iungman, T., Cirach, M., Marando, F., Barboza, E.P., Khomenko, S., Masselot, P., Quijal-Zamorano, M., Mueller, N., Gasparrini, A., Urquiza, J., Heris, M., Thondoo, M., Nieuwenhuijsen, M., 2023. Cooling cities through urban green infrastructure: a health impact assessment of European cities. The Lancet 401, 577–589. 10.1016/S0140-6736(22)02585-5

Jepson, V., Apparicio, P., Pham, T.-T.-H., 2022. Environmental equity and access to parks in Greater Montreal: an analysis of spatial proximity and potential congestion issues. Journal of Urbanism 1–19.

Jolivet, V., Carré, M.-N., 2017. Métabolisme urbain et quartiers péricentraux dans la métropolisation. L’exemple du quartier de Saint-Michel à Montréal. Cybergeo: European Journal of Geography. 10.4000/cybergeo.28067

Kay, M., Mastny, T., 2023. tidybayes: Tidy Data and “Geoms” for Bayesian Models.

Kiani, B., Thierry, B., Fuller, D., Firth, C., Winters, M., Kestens, Y., 2023. Gentrification, neighborhood socioeconomic factors and urban vegetation inequities: A study of greenspace and tree canopy increases in Montreal, Canada. Landscape and Urban Planning 240, 104871. 10.1016/j.landurbplan.2023.104871

Kikano, F., Kingsbury, C., Landry-Duval, M.-A., Fréchette, L., 2023. Portrait de la Gouvernance des Parcs À Montréal: Les Cas du Square Cabot, de la Place Émilie-Gamelin et des Parcs Jarry et Frédéric-Back (Revue de la littérature). Centre d’étude en responsabilité sociale et écocitoyenneté (CÉRSÉ).

Koebel, C.T., 1996. Urban Redevelopment, Displacement and the Future of the American City (Technical Report). Virginia Center for Housing Research.

Kozlowski, T.T., 1999. Soil Compaction and Growth of Woody Plants. Scandinavian Journal of Forest Research 14, 596–619. 10.1080/02827589908540825

Kraemer, R., Kabisch, N., 2022. Parks Under Stress: Air Temperature Regulation of Urban Green Spaces Under Conditions of Drought and Summer Heat. Frontiers in Environmental Science 10.

Kremer, P., Hamstead, Z., Haase, D., McPhearson, T., Frantzeskaki, N., Andersson, E., Kabisch, N., Larondelle, N., Rall, E.L., Voigt, A., Baró, F., Bertram, C., Gómez-Baggethun, E., Hansen, R., Kaczorowska, A., Kain, J.-H., Kronenberg, J., Langemeyer, J., Pauleit, S., Rehdanz, K., Schewenius, M., van Ham, C., Wurster, D., Elmqvist, T., 2016. Key insights for the future of urban ecosystem services research. Ecology and Society 21, 1–11.

Landry, F., Dupras, J., Messier, C., 2020. Convergence of urban forest and socio-economic indicators of resilience: A study of environmental inequality in four major cities in eastern Canada.

Landscape and Urban Planning 202, 103856. 10.1016/j.landurbplan.2020.103856

Larsen, K., Gilliland, J., 2008. Mapping the evolution of “food deserts” in a Canadian city: Supermarket accessibility in London, Ontario, 1961–2005. International Journal of Health Geographics 7, 16. 10.1186/1476-072X-7-16

Little, E.L., 1980. National Audubon Society Field Guide to Trees: Eastern Region, North America. Alfred A Knopf, New York.

Magarik, Y.A.S., Roman, L.A., Henning, J.G., 2020. How should we measure the DBH of multi-stemmed urban trees? Urban Forestry & Urban Greening 47, 126481. 10.1016/j.ufug.2019.126481

Maynard, R., 2017. Policing Black lives: State violence in Canada from slavery to the present. Fernwood Publishing.

McElreath, R., 2020. Statistical rethinking: a Bayesian course with examples in R and Stan, Second edition. ed, Chapman & Hall/CRC texts in statistical science series. CRC Press, Boca Raton London New York.

McLauchlan, K., 2006. The Nature and Longevity of Agricultural Impacts on Soil Carbon and Nutrients: A Review. Ecosystems 9, 1364–1382.

McPherson, E.G., Nowak, D., Heisler, G., Grimmond, S., Souch, C., Grant, R., Rowntree, R., 1997. Quantifying urban forest structure, function, and value: the Chicago Urban Forest Climate Project. Urban Ecosystems 1, 49–61. 10.1023/A:1014350822458

Mennis, J., 2006. Socioeconomic-Vegetation Relationships in Urban, Residential Land. Photogrammetric Engineering & Remote Sensing 72, 911–921. 10.14358/PERS.72.8.911

Mills, B.R., 2022. MetBrewer: Color Palettes Inspired by Works at the Metropolitan Museum of Art.

Munteanu, C., Kuemmerle, T., Keuler, N.S., Müller, D., Balázs, P., Dobosz, M., Griffiths, P., Halada, L., Kaim, D., Király, G., Konkoly-Gyuró, É., Kozak, J., Lieskovsky, J., Ostafin, K., Ostapowicz, K., Shandra, O., Radeloff, V.C., 2015. Legacies of 19th century land use shape contemporary forest cover. Global Environmental Change 34, 83–94. 10.1016/j.gloenvcha.2015.06.015

Ndiaye, S., 2004. Projet de recherche sur les quinze ans de la table de concertation : Vivre Saint-Michel en Santé (VSMS) Phase I : Profil de la communauté (No. R-16-2004). Centraide du Grand Montréal.

Nelson, S.H., Bigger, P., 2022. Infrastructural nature. Progress in Human Geography 46, 86–107. 10.1177/0309132521993916

Nix, S., Roman, L.A., Healy, M., Rogan, J., Pearsall, H., 2023. Linking tree cover change to historical management practices in urban parks. Landsc Ecol 38, 4227–4245. 10.1007/s10980-022-01543-4

Ossola, A., Cadenasso, M.L., Meineke, E.K., 2021. Valuing the Role of Time in Urban Ecology. Front. Ecol. Evol. 9, 620620. 10.3389/fevo.2021.620620

Ossola, A., Locke, D., Lin, B., Minor, E., 2019. Greening in style: Urban form, architecture and the structure of front and backyard vegetation. Landscape and Urban Planning 185, 141–157. 10.1016/j.landurbplan.2019.02.014

Pabst, H., Gerschlauer, F., Kiese, R., Kuzyakov, Y., 2016. Land Use and Precipitation Affect Organic and Microbial Carbon Stocks and the Specific Metabolic Quotient in Soils of Eleven Ecosystems of Mt. Kilimanjaro, Tanzania. Land Degradation & Development 27, 592–602. 10.1002/ldr.2406

Padgham, M., Rudis, B., Lovelace, R., Salmon, M., Maspons, J., Smith, A., Smith, J., Gilardi, A., Spinielli, E., North, A., Machyna, M., code), M.K. (Author of included R., Pousson, E., 2023. osmdata: Import “OpenStreetMap” Data as Simple Features or Spatial Objects.

Pallagst, K., 2009. Shrinking Cities in the United States of America, in: The Future of Shrinking Cities - Problems, Patterns and Strategies of Urban Transformation in a Global Context. Institute of Urban and Regional Development, Center for Global Metropolitan Studies, and the Shrinking Cities International Research Network Monograph Series, pp. 81–88.

Parish, J., 2020. Re-wilding Parkdale? Environmental gentrification, settler colonialism, and the reconfiguration of nature in 21st century Toronto. Environment and Planning E: Nature and Space 3, 263–286. 10.1177/2514848619868110

Pebesma, E., Bivand, R., Racine, E., Sumner, M., Cook, I., Keitt, T., Lovelace, R., Wickham, H., Ooms, J., Müller, K., Pedersen, T.L., Baston, D., Dunnington, D., 2023a. sf: Simple Features for R.

Pebesma, E., Sumner, M., Racine, E., Fantini, A., Blodgett, D., 2023b. stars: Spatiotemporal Arrays, Raster and Vector Data Cubes.

Pham, T.-T.-H., Apparicio, P., Landry, S., Séguin, A.-M., Gagnon, M., 2013. Predictors of the distribution of street and backyard vegetation in Montreal, Canada. Urban Forestry & Urban Greening 12, 18–27. 10.1016/j.ufug.2012.09.002

Pham, T.-T.-H., Apparicio, P., Séguin, A.-M., Landry, S., Gagnon, M., 2012. Spatial distribution of vegetation in Montreal: An uneven distribution or environmental inequity? Landscape and Urban Planning 107, 214–224. 10.1016/j.landurbplan.2012.06.002

Powell, J.A., 2008. Structural racism: Building upon the insights of John Calmore. North Carolina Law Review 86, 26.

Pregitzer, C.C., Bradford, M.A., 2023. Associations between recent land use history and urban forest composition. Urban Forestry & Urban Greening 90, 128134. 10.1016/j.ufug.2023.128134

QGIS Development Team, 2020. QGIS Geographic Information System. Hannover.

Qiu, J., Carpenter, S.R., Booth, E.G., Motew, M., Zipper, S.C., Kucharik, C.J., Loheide II, S.P., Turner, M.G., 2018. Understanding relationships among ecosystem services across spatial scales and over time. Environmental Research Letters 13, 1–16.

R Core Team, 2023. R: A Language and Environment for Statistical Computing.

Raciti, S.M., Groffman, P.M., Jenkins, J.C., Pouyat, R.V., Fahey, T.J., Pickett, S.T.A., Cadenasso, M.L., 2011. Accumulation of Carbon and Nitrogen in Residential Soils with Different Land-Use Histories. Ecosystems 14, 287–297. 10.1007/s10021-010-9409-3

Rahman, M.A., Stratopoulos, L.M.F., Moser-Reischl, A., Zölch, T., Häberle, K.-H., Rötzer, T., Pretzsch, H., Pauleit, S., 2020. Traits of trees for cooling urban heat islands: A meta-analysis. Building and Environment 170, 106606. 10.1016/j.buildenv.2019.106606

Remme, R.P., Frumkin, H., Guerry, A.D., King, A.C., Mandle, L., Sarabu, C., Bratman, G.N., Giles-Corti, B., Hamel, P., Han, B., Hicks, J.L., James, P., Lawler, J.J., Lindahl, T., Liu, H., Lu, Y., Oosterbroek, B., Paudel, B., Sallis, J.F., Schipperijn, J., Sosič, R., de Vries, S., Wheeler, B.W., Wood, S.A., Wu, T., Daily, G.C., 2021. An ecosystem service perspective on urban nature, physical activity, and health. Proc Natl Acad Sci USA 118, e2018472118. 10.1073/pnas.2018472118

Rivière-des-Prairies–Pointe-aux-Trembles, 2013. Arbres: entretien, abattage, protection, plantation et aménagement paysager des terrains. Ville de Montréal.

Robillard, A., Boisjoly, G., Waygood, E.O.D., 2023. Access to Parks and Green Spaces in Quebec City, Canada: Developing Children-Specific Accessibility Measures. Transportation Research Record 2677, 464–477. 10.1177/03611981231161618

Robinson, G.R., Handel, S.N., Schmalhofer, V.R., 1992. Survival, reproduction, and recruitment of woody plants after 14 years on a reforested landfill. Environmental Management 16, 265–271. 10.1007/BF02393832

Roman, L., Catton, I., Greenfield, E., Pearsall, H., Eisenman, T., Henning, J., 2021. Linking Urban Tree Cover Change and Local History in a Post-Industrial City. Land 10, 403. 10.3390/land10040403

Roman, L.A., Pearsall, H., Eisenman, T.S., Conway, T.M., Fahey, R.T., Landry, S., Vogt, J., van Doorn, N.S., Grove, J.M., Locke, D.H., Bardekjian, A.C., Battles, J.J., Cadenasso, M.L., van den Bosch, C.C.K., Avolio, M., Berland, A., Jenerette, G.D., Mincey, S.K., Pataki, D.E., Staudhammer, C., 2018. Human and biophysical legacies shape contemporary urban forests: A literature synthesis. Urban Forestry & Urban Greening 31, 157–168. 10.1016/j.ufug.2018.03.004

Rooney, R.C., Bayley, S.E., Schindler, D.W., 2012. Oil sands mining and reclamation cause massive loss of peatland and stored carbon. Proceedings of the National Academy of Sciences 109, 4933–4937. 10.1073/pnas.1117693108

Saint-Laurent, 2018. Arbres: plantation et aménagement paysager. Ville de Montréal.

Sarofim, M.C., Saha, S., Hawkins, M.D., Mills, D.M., Hess, J., Horton, R., Kinney, P., Schwartz, J., St. Juliana, A., 2016. Ch. 2: Temperature-Related Death and Illness. The Impacts of Climate Change on Human Health in the United States: A Scientific Assessment. U.S. Global Change Research Program. 10.7930/J0MG7MDX

Sax, D.L., Nesbitt, L., Quinton, J., 2022. Improvement, not displacement: A framework for urban green gentrification research and practice. Environmental Science & Policy 137, 373–383. 10.1016/j.envsci.2022.09.013

Schell, C.J., Dyson, K., Fuentes, T.L., Des Roches, S., Harris, N.C., Miller, D.S., Woelfle-Erskine, C.A., Lambert, M.R., 2020. The ecological and evolutionary consequences of systemic racism in urban environments. Science 369. 10.1126/science.aay4497

Shah, T.I., Bell, S., Wilson, K., 2016. Spatial Accessibility to Health Care Services: Identifying under-Serviced Neighbourhoods in Canadian Urban Areas. PLoS ONE 11, e0168208. 10.1371/journal.pone.0168208

Shumi, G., Schultner, J., Dorresteijn, I., Rodrigues, P., Hanspach, J., Hylander, K., Senbeta, F., Fischer, J., 2018. Land use legacy effects on woody vegetation in agricultural landscapes of south-western Ethiopia. Diversity and Distributions 24, 1136–1148. 10.1111/ddi.12754

Soto, J.R., Escobedo, F.J., Khachatryan, H., Adams, D.C., 2018. Consumer demand for urban forest ecosystem services and disservices: Examining trade-offs using choice experiments and best-worst scaling. Ecosystem Services 29, 31–39. 10.1016/j.ecoser.2017.11.009

Statistics Canada, 2022. Census Profile 2021 Census.

Stickley, S.F., Fraterrigo, J.M., 2021. Understory vegetation contributes to microclimatic buffering of near-surface temperatures in temperate deciduous forests. Landscape Ecol 36, 1197–1213. 10.1007/s10980-021-01195-w

Tappeiner, U., Leitinger, G., Zariņa, A., Bürgi, M., 2020. How to consider history in landscape ecology: patterns, processes, and pathways. Landscape Ecol. 10.1007/s10980-020-01163-w

Thieurmel, B., Elmarhraoui, A., 2022. suncalc: Compute Sun Position, Sunlight Phases, Moon Position and Lunar Phase.

Thompson, J., Brokaw, N., Zimmerman, J.K., Waide, R.B., Everham III, E.M., Lodge, D.J., Taylor, C.M., García-Montiel, D., Fluet, M., 2002. Land Use History, Environment, and Tree Composition in a Tropical Forest. Ecological Applications 12, 1344–1363. 10.1890/1051-0761(2002)012[1344:LUHEAT]2.0.CO;2

Triguero-Mas, M., Anguelovski, I., García-Lamarca, M., Argüelles, L., Perez-del-Pulgar, C., Shokry, G., Connolly, J.J.T., Cole, H.V.S., 2021. Natural outdoor environments’ health effects in gentrifying neighborhoods: Disruptive green landscapes for underprivileged neighborhood residents. Social Science & Medicine 279, 113964. 10.1016/j.socscimed.2021.113964

Ture, K., Hamilton, C.V., 1992. Black power: The politics of liberation in America. Vintage Books.

United Church of Christ, 1987. Toxic Wastes and Race In the United States: A National Report on the Racial and Socio-Economic Characteristics of Communities with Hazardous Waste Sites. Commission for Racial Justice.

Vaz Monteiro, M., Doick, K.J., Handley, P., Peace, A., 2016. The impact of greenspace size on the extent of local nocturnal air temperature cooling in London. Urban Forestry & Urban Greening 16, 160–169. 10.1016/j.ufug.2016.02.008

Villamagna, A.M., Angermeier, P.L., Bennett, E.M., 2013. Capacity, pressure, demand, and flow: A conceptual framework for analyzing ecosystem service provision and delivery. Ecological Complexity 15, 114–121. 10.1016/j.ecocom.2013.07.004

Ville de Montréal, 2023a. Boroughs [WWW Document]. URL https://montreal.ca/en/boroughs (accessed 10.17.23).

Ville de Montréal, 2023b. Parcours riverain.

Waldron, I., 2018. Re-thinking waste: mapping racial geographies of violence on the colonial landscape. Environmental Sociology 4, 36–53.

Watkins, S.L., Gerrish, E., 2018. The relationship between urban forests and race: A meta-analysis. J Environ Manage 209, 152–168. 10.1016/j.jenvman.2017.12.021

Wickham, H., Chang, W., Henry, L., Pedersen, T.L., Takahashi, K., Wilke, C., Woo, K., Yutani, H., Dunnington, D., Posit, PBC, 2023. ggplot2: Create Elegant Data Visualisations Using the Grammar of Graphics.

World Meteorological Association, 2021. Guide to Instruments and Methods of Observation [2021 edition - Volume I: Measurement of Meteorological Variables] (No. WMO-No. 8). Geneva, Switzerland.

Wujeska-Klause, A., Pfautsch, S., 2020. The Best Urban Trees for Daytime Cooling Leave Nights Slightly Warmer. Forests 11, 945. 10.3390/f11090945

Yang, J., Yan, P., He, R., Song, X., 2017. Exploring land-use legacy effects on taxonomic and functional diversity of woody plants in a rapidly urbanizing landscape. Landscape and Urban Planning 162, 92–103. 10.1016/j.landurbplan.2017.02.003

Zhao, J., Zhao, X., Wu, D., Meili, N., Fatichi, S., 2023. Satellite-based evidence highlights a considerable increase of urban tree cooling benefits from 2000 to 2015. Global Change Biology 29, 3085–3097.

Ziter, C., Bennett, E.M., Gonzalez, A., 2013. Functional diversity and management mediate aboveground carbon stocks in small forest fragments. Ecosphere 4, 1–21. 10.1890/ES13-00135.1

Ziter, C., Graves, R.A., Turner, M.G., 2017. How do land-use legacies affect ecosystem services in United States cultural landscapes? Landscape Ecol 32, 2205–2218. 10.1007/s10980-017-0545-4

Ziter, C., Turner, M.G., 2018. Current and historical land use influence soil-based ecosystem services in an urban landscape. Ecological Applications 28, 643–654. 10.1002/eap.1689

Ziter, C.D., Pedersen, E.J., Kucharik, C.J., Turner, M.G., 2019. Scale-dependent interactions between tree canopy cover and impervious surfaces reduce daytime urban heat during summer. Proc Natl Acad Sci USA 116, 7575–7580. 10.1073/pnas.1817561116

Zwolak, A., Sarzyńska, M., Szpyrka, E., Stawarczyk, K., 2019. Sources of Soil Pollution by Heavy Metals and Their Accumulation in Vegetables: a Review. Water Air Soil Pollut 230, 1–9. 10.1007/s11270-019-4221-y

