## Appendix S1 for "Land-use history causes differences in park nighttime cooling capacity and forest structure"

### Figures

#### Figure S1

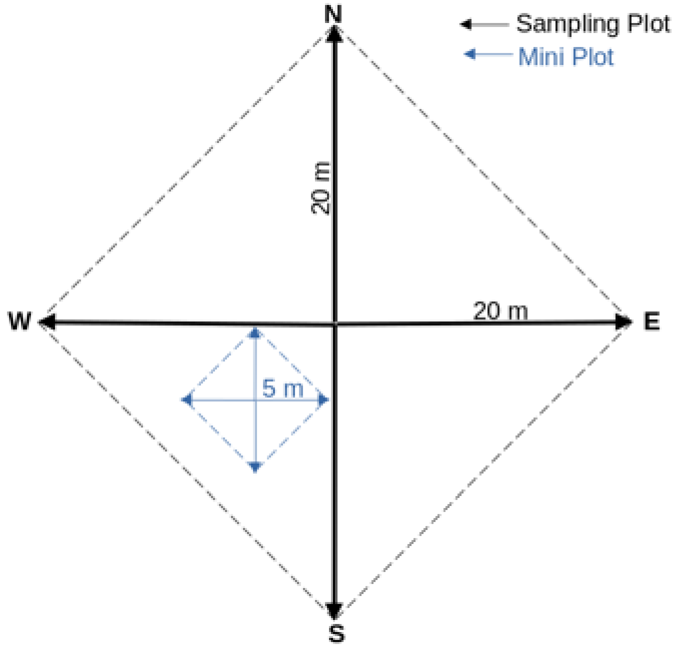

Figure S1. Sampling plot configuration. Mini-plot was located in the most representative part of the larger plot, as decided in the field. Mini-Plot Information: due to the nature of the permits we received, most of the "full plot" sampling of trees < 5 cm was done in typical urban parks (n = 11/14 parks). These parks are characterized by large grassy lawns, with a low density of very large deciduous trees. There are sometimes new plantings which results in small trees, but very few of them. For many of these plots, there were no trees < 5 cm even when the "full plot" is used. We switched to the mini plot method when we started sampling "nature parks", where there is a more natural forest ecosystem. In these systems, we could find hundreds of trees < 5 cm, even within our mini plot. Thus, the full plots are very similar or identical to what the mini plots would look like in the majority of cases, as they either have 0 or 1 trees < 5 cm.

### Figure S2

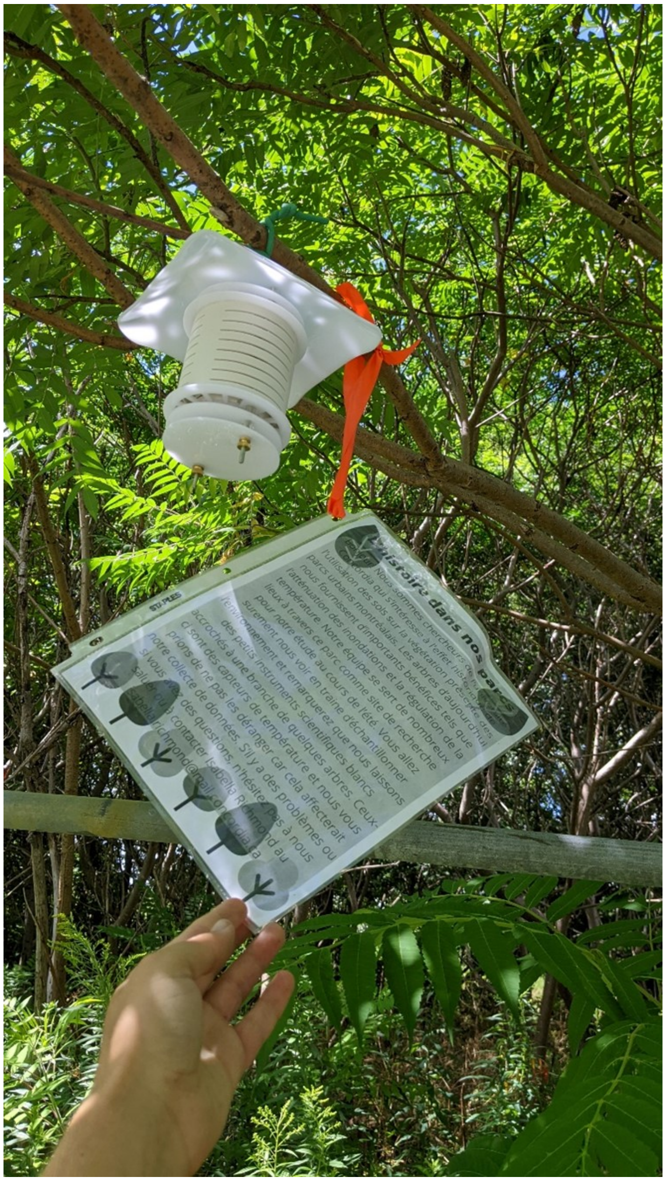

Figure S2. Example sensor setup. Sensor is a CredoSense CSL-T0.5 temperature logger located within a 3-D printed shield and hung with an informational flyer containing the contact information of the lead author. Photo taken by Isabella C Richmond.

### Figure S3

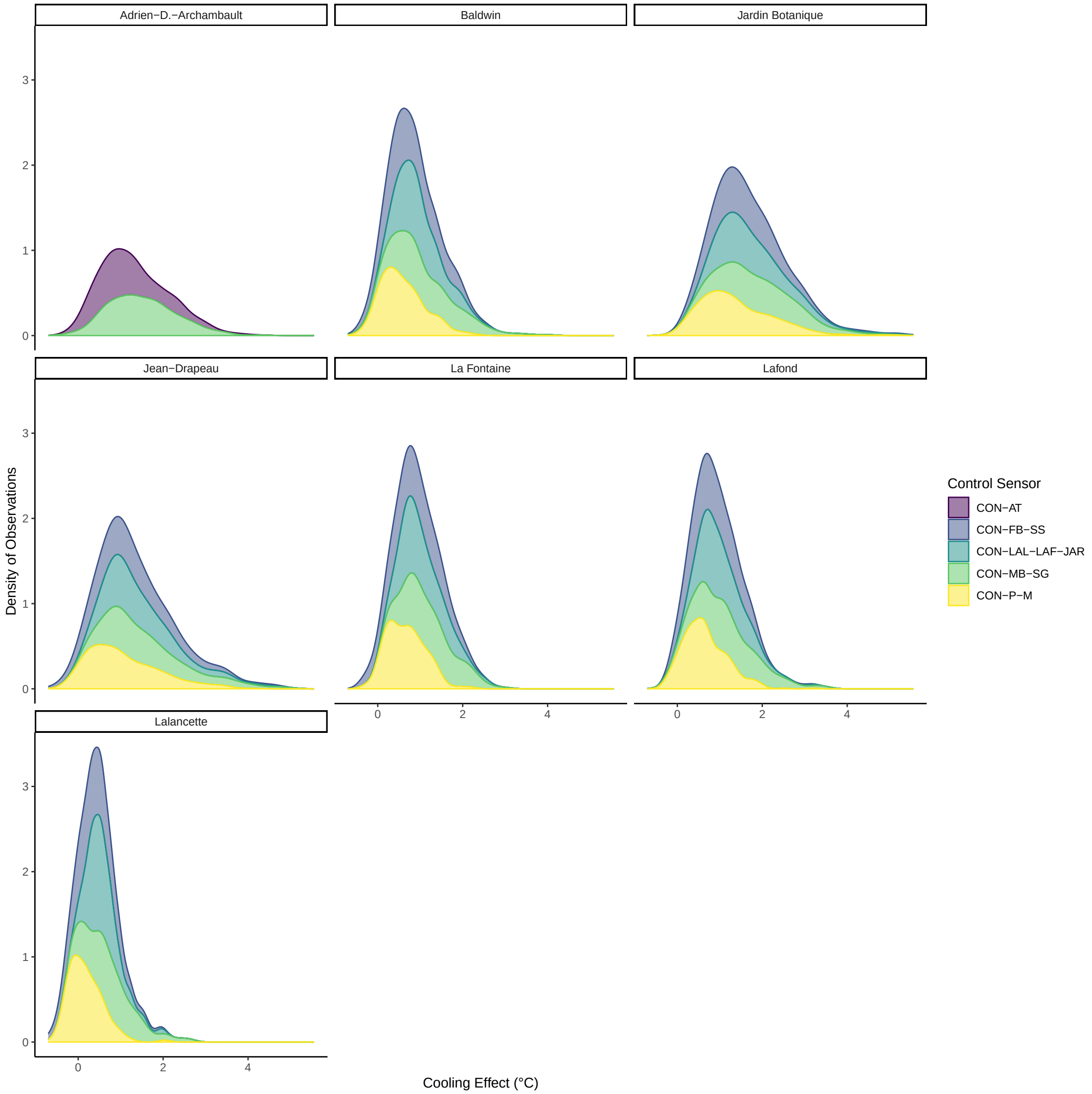

Figure S3. Sensitivity analysis showing the cooling effect across all dates for each park where the control sensor was replaced due to missing data. Plots were made with the remaining control sensors closest to the park in question. CON-LAL-LAF-JAR indicates the control sensor that we used from another, similar study.

### Figure S4

Below are the math stats notations for all models run in our paper.

#### Total Effect of Past Land-Use on Cooling Effect

$$
\begin{aligned}
&Cooling_i \sim Normal(\mu_i, \sigma)\\
&\mu_i \sim \alpha_{Park[i]} + \alpha_{Plot[i]} + \gamma_{PLU[i]} + \gamma_{tod[i]} + \beta_{PLU}tod + \beta_{PLU[i]} + \beta_{tod[i]} + \beta_{age}age_i\\
&\alpha_{Park[j]} \sim Normal(\bar{\alpha}, \sigma_{\alpha_{Park}})\\
&\alpha_{Plot[j]} \sim Normal(0, \sigma_{\alpha_{Plot}})\\
&\gamma_j \sim Normal(0, 0.5)\\
&\beta_j \sim Normal(0, 0.5)\\
&\bar{\alpha} \sim Normal(0, 0.5)\\
&\sigma_{\alpha_{Park}} \sim Half-Normal(0, 0.2)\\
&\sigma_{\alpha_{Plot}} \sim Half-Normal(0, 0.2)\\
&\sigma \sim Exponential(1)
\end{aligned}
$$

- $\alpha$ and $\gamma$ both represent intercepts, $\alpha$ is used for random effects and $\gamma$ for fixed interaction effects
- $Cooling_i$ is the centered and scaled cooling effect in each plot in degrees Celsuis
- $Park_i$ is the park that each plot is in
- $Plot_i$ is the plot that each measurement is in
- $PLU_i$ is the past land use category that each measurement is in (i.e., agricultural, forested, industrial)
- $tod_i$ is the time of day category that each measurement is in (i.e., daytime, nighttime)
- $age_i$ is the centered and scaled age of the park that each measurement is in

#### Direct Effect of Past Land-Use on Forest Composition

#### Large Tree Density

$$
\begin{aligned}
&DensL_i \sim Normal(\mu_i, \sigma)\\
&\mu_i = \alpha + \beta_1 forested_{[i]} + \beta_2 industrial_{[i]}\\
&\alpha \sim Normal(0, 0.5)\\
&\beta_j \sim Normal(0, 0.5)\\
&\sigma \sim Exponential(1)
\end{aligned}
$$

Where:

- $DensL_i$ is the centered and scaled basal area of large trees in each plot
- `forested` is the factor level from `PastLandUse` that indicates previously forested parks. Previously agricultural parks are the default level, absorbed into the intercept.
- `industrial` is the factor level from `PastLandUse` that indicates previously forested parks. Previously agricultural parks are the default level, absorbed into the intercept.

##### Small Tree Density

$$
\begin{aligned}
&log(DensS_i) \sim Normal(\mu_i, \sigma)\\
&\mu_i = \alpha + \beta_1 forested_{[i]} + \beta_2 industrial_{[i]}\\
&\alpha \sim Normal(0, 0.5)\\
&\beta_j \sim Normal(0, 0.5)\\
&\sigma \sim Exponential(1)
\end{aligned}
$$

- $log(DensS_i)$ is the centered and scaled density of small trees in each plot. Logged for model fit.
- `forested` is the factor level from `PastLandUse` that indicates previously forested parks. Previously agricultural parks are the default level, absorbed into the intercept.
- `industrial` is the factor level from `PastLandUse` that indicates previously forested parks. Previously agricultural parks are the default level, absorbed into the intercept.

##### Large Tree Size

$$
\begin{aligned}
&SizeL_i \sim Normal(\mu_i, \sigma)\\
&\mu_i = \alpha + \beta_1 forested_{[i]} + \beta_2 industrial_{[i]}\\
&\alpha \sim Normal(-1, 0.5)\\
&\beta_j \sim Normal(0, 0.5)\\
&\sigma \sim Exponential(1)
\end{aligned}
$$

- $SizeL_i$ is the centered and scaled median size of large trees in each plot.
- `forested` is the factor level from `PastLandUse` that indicates previously forested parks. Previously agricultural parks are the default level, absorbed into the intercept.
- `industrial` is the factor level from `PastLandUse` that indicates previously forested parks. Previously agricultural parks are the default level, absorbed into the intercept.

#### Small Tree Size

$$
\begin{aligned}
SizeS_i &\sim Normal(\mu_i, \sigma) \\
\mu_i &= \alpha + \beta_1 forested_{[i]} + \beta_2 industrial_{[i]} \\
\alpha &\sim Normal(1, 0.5) \\
\beta_j &\sim Normal(0, 0.5) \\
\sigma &\sim Exponential(1)
\end{aligned}
$$

- $SizeS_i$ is the centered and scaled median size of small trees in each plot.
- `forested` is the factor level from `PastLandUse` that indicates previously forested parks. Previously agricultural parks are the default level, absorbed into the intercept.
- `industrial` is the factor level from `PastLandUse` that indicates previously forested parks. Previously agricultural parks are the default level, absorbed into the intercept.

#### Large Tree Species Richness

$$
\begin{aligned}
SRL_i &\sim Normal(\mu_i, \sigma) \\
\mu_i &= \alpha + \beta_1 forested_{[i]} + \beta_2 industrial_{[i]} \\
\alpha &\sim Normal(2, 1) \\
\beta_j &\sim Normal(0, 0.5) \\
\sigma &\sim Exponential(1)
\end{aligned}
$$

- $SRL_i$ is the centered and scaled species richness of large trees in each plot.
- `forested` is the factor level from `PastLandUse` that indicates previously forested parks. Previously agricultural parks are the default level, absorbed into the intercept.
- `industrial` is the factor level from `PastLandUse` that indicates previously forested parks. Previously agricultural parks are the default level, absorbed into the intercept.

#### Direct Effect of Forest Composition on Cooling Effect

$$Cooling_i \sim Normal(\mu_i, \sigma)$$
$$\mu_i \sim \alpha_{Park[i]} + \alpha_{Plot[i]} + \gamma_{tod[i]} + \beta_{tod[i]}DensityL_i + \beta_{tod[i]}DensityS_i + \beta_{tod[i]}SizeL_i$$
$$+ \beta_{tod[i]}SizeS_i + \beta_{tod[i]}SRL_i + \beta_{DensityL}DensityL_i + \beta_{DensityS}DensityS_i + \beta_{SizeL}SizeL_i$$
$$+ \beta_{SizeS}SizeS_i + \beta_{SRL}SRL_i$$
$$\alpha_{Park[j]} \sim Normal(\bar{\alpha}, \sigma_{\alpha_{Park}})$$
$$\alpha_{Plot[j]} \sim Normal(0, \sigma_{\alpha_{Plot}})$$
$$\gamma_j \sim Normal(0, 0.5)$$
$$\beta_j \sim Normal(0, 0.5)$$
$$\bar{\alpha} \sim Normal(0, 0.5)$$
$$\sigma_{\alpha_{Park}} \sim Half - Normal(0, 0.2)$$
$$\sigma_{\alpha_{Plot}} \sim Half - Normal(0, 0.2)$$
$$\sigma \sim Exponential(1)$$

- $\alpha$ and $\gamma$ both represent intercepts, $\alpha$ is used for random effects and $\gamma$ for fixed interaction effecst
- $Cooling_i$ is the centered and scaled cooling effect in each plot in degrees Celsuis
- $Park_i$ is the park that each plot is in
- $Plot_i$ is the plot that each measurement is in
- $PLU_i$ is the past land use category that each measurement is in (i.e., agricultural, forested, industrial)
- $tod_i$ is the time of day category that each measurement is in (i.e., daytime, nighttime)
- $age_i$ is the centered and scaled age of the park that each measurement is in

### Figure S5

#### Prior Predictive Checks

Prior predictive checks are used to ensure that the values selected for priors for our models allow a biologically reasonable range of values. For models with fixed effects fit to the model (models 1 and 5), we simulate predictive draws for prior only models and visualize the slope/intercept of the values. We then do a "posterior predictive check" but with the prior only model, to see if the data is captured in the priors. For models with only random effects, we only use the posterior predictive check.

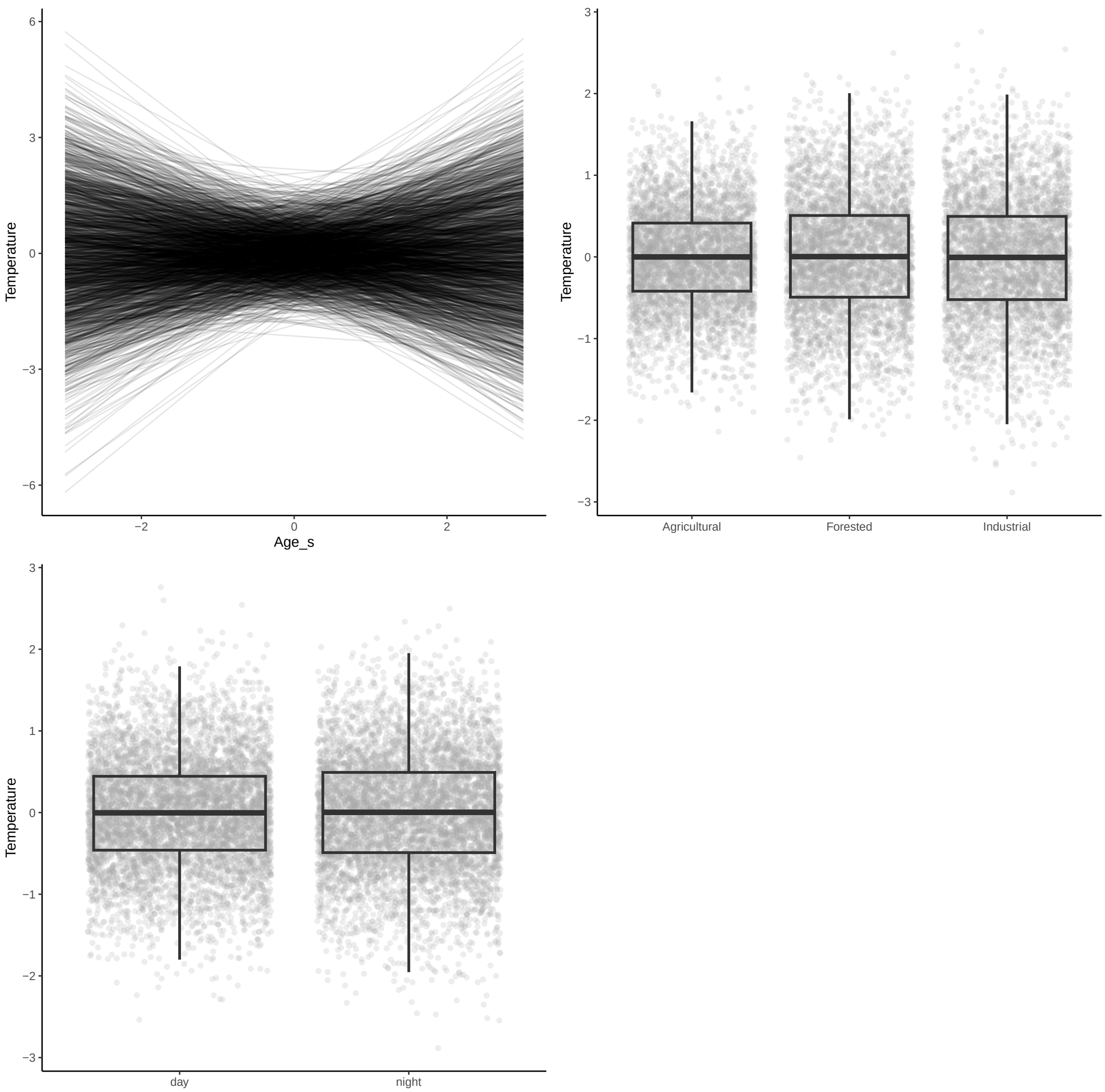

Figure S5a. Model 1 - Total Effect of Past Land-Use on Temperature

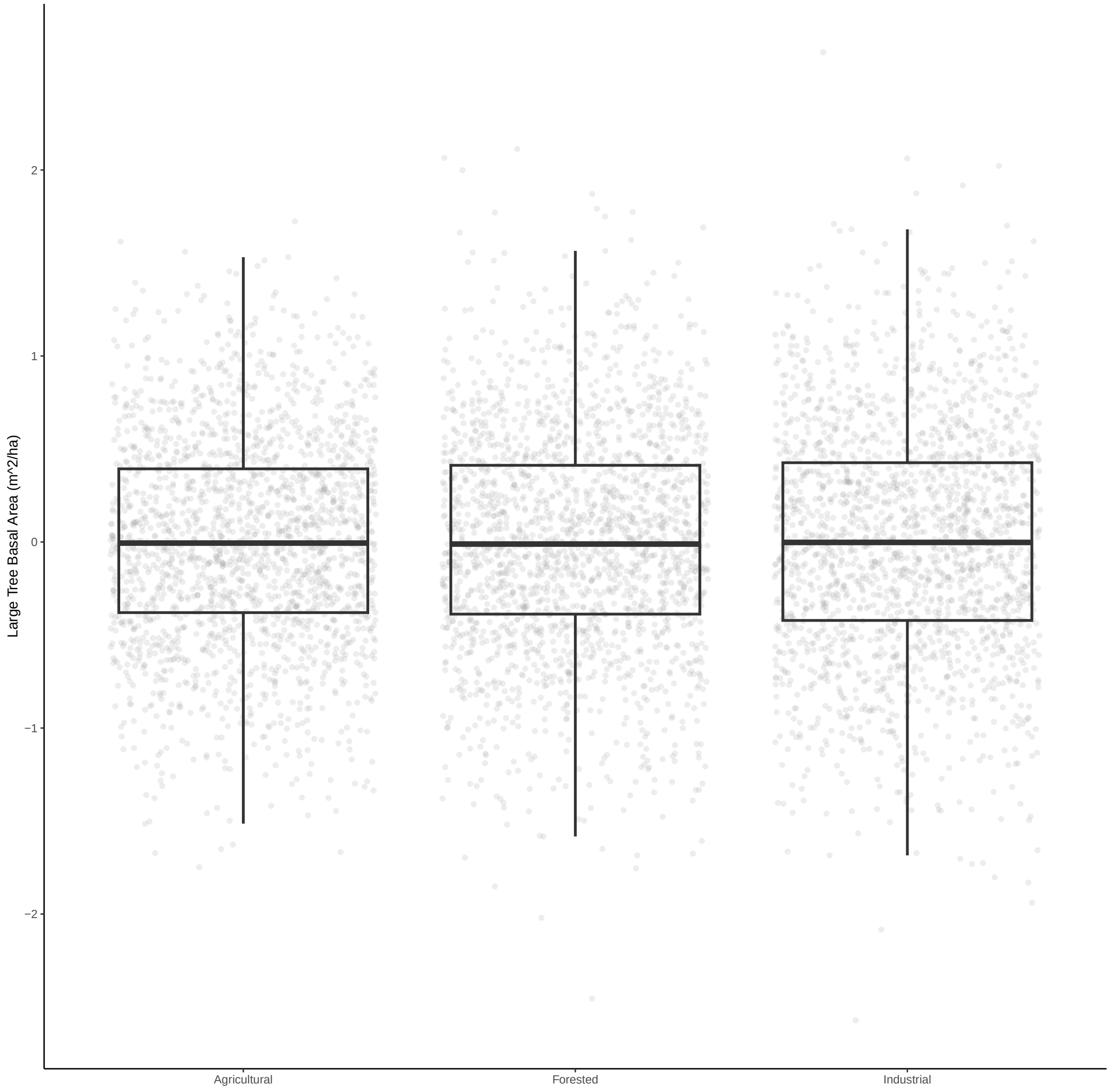

Figure S5b. Model 2a - Direct Effect of Past Land-Use on Large Tree (>= 5 cm DBH) Basal Area

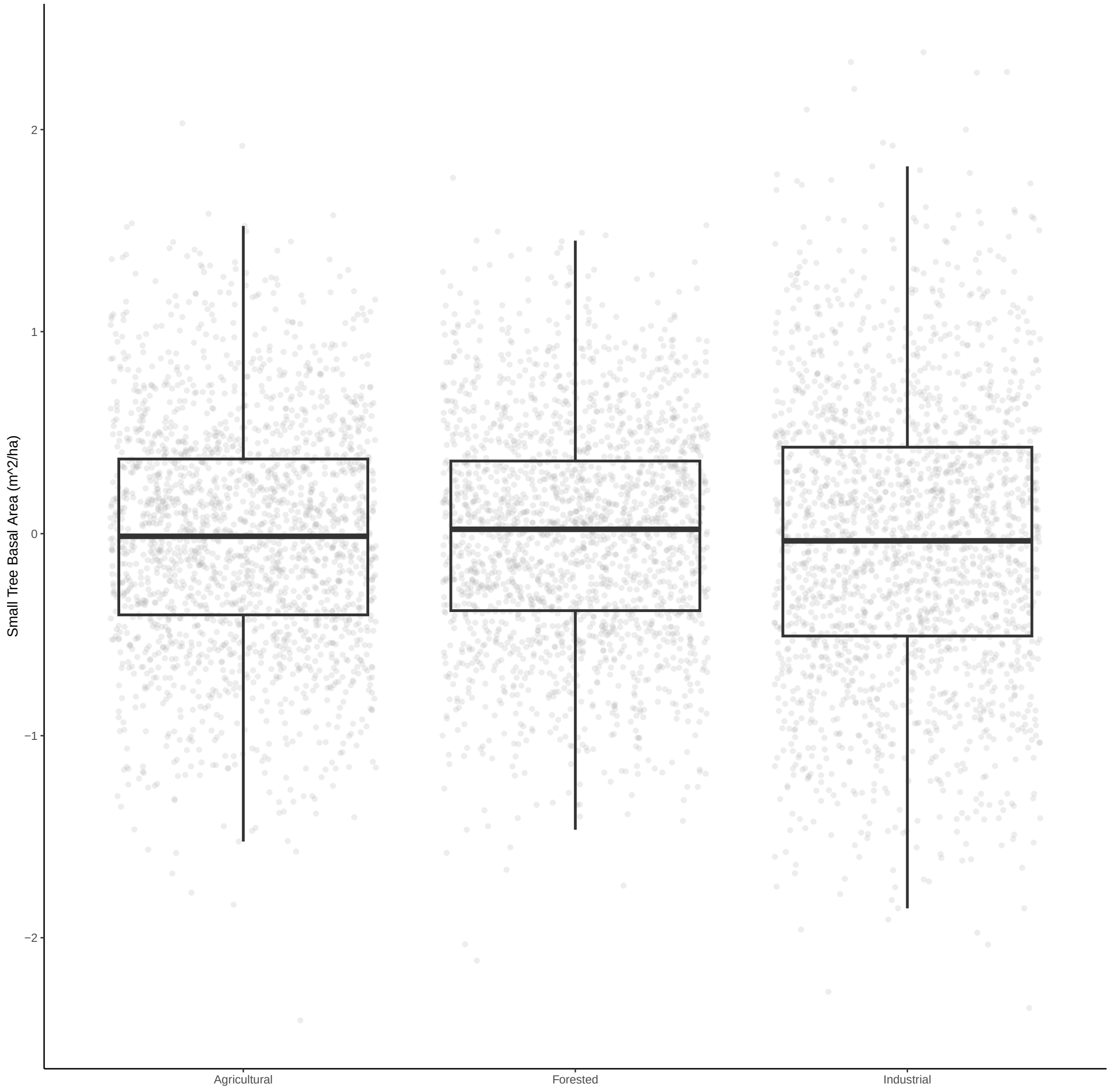

Figure S5c. Model 2b - Direct Effect of Past Land-Use on Small Tree (< 5 cm DBH) Density

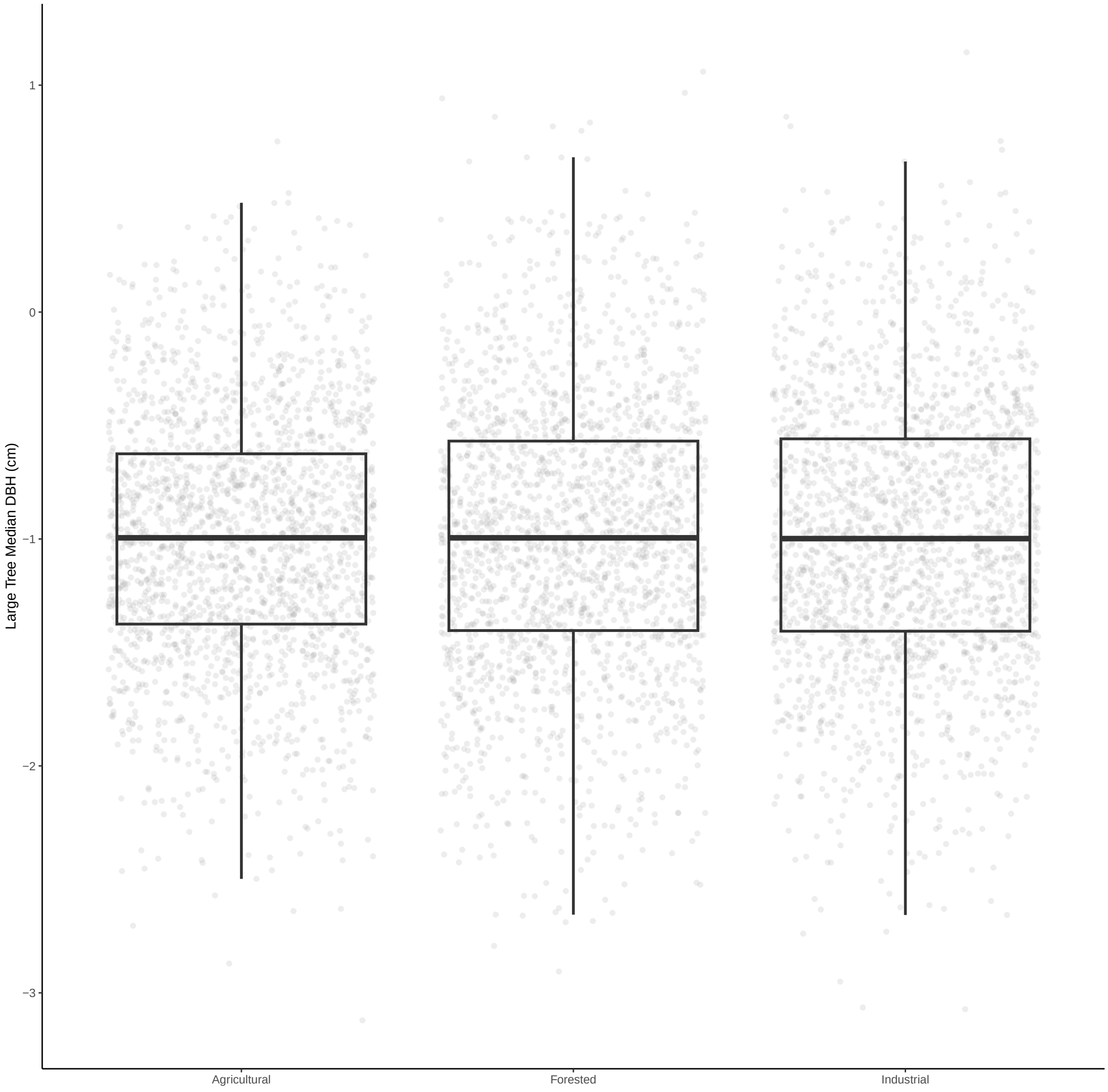

Figure S5d. Model 3a - Direct Effect of Past Land-Use on Large Tree (>= 5 cm DBH) Size

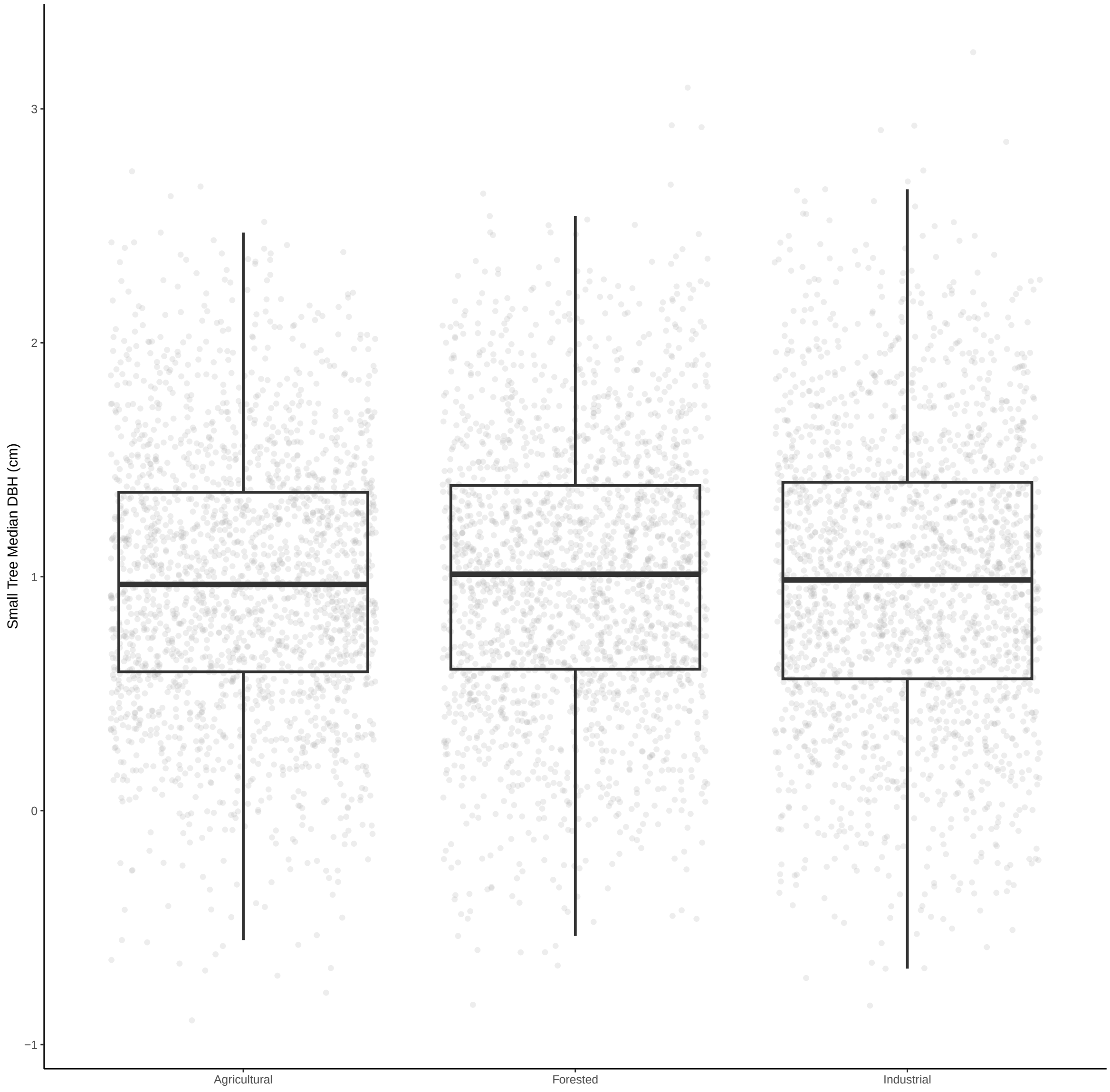

Figure S5e. Model 3b - Direct Effect of Past Land-Use on Small Tree (< 5 cm DBH) Size

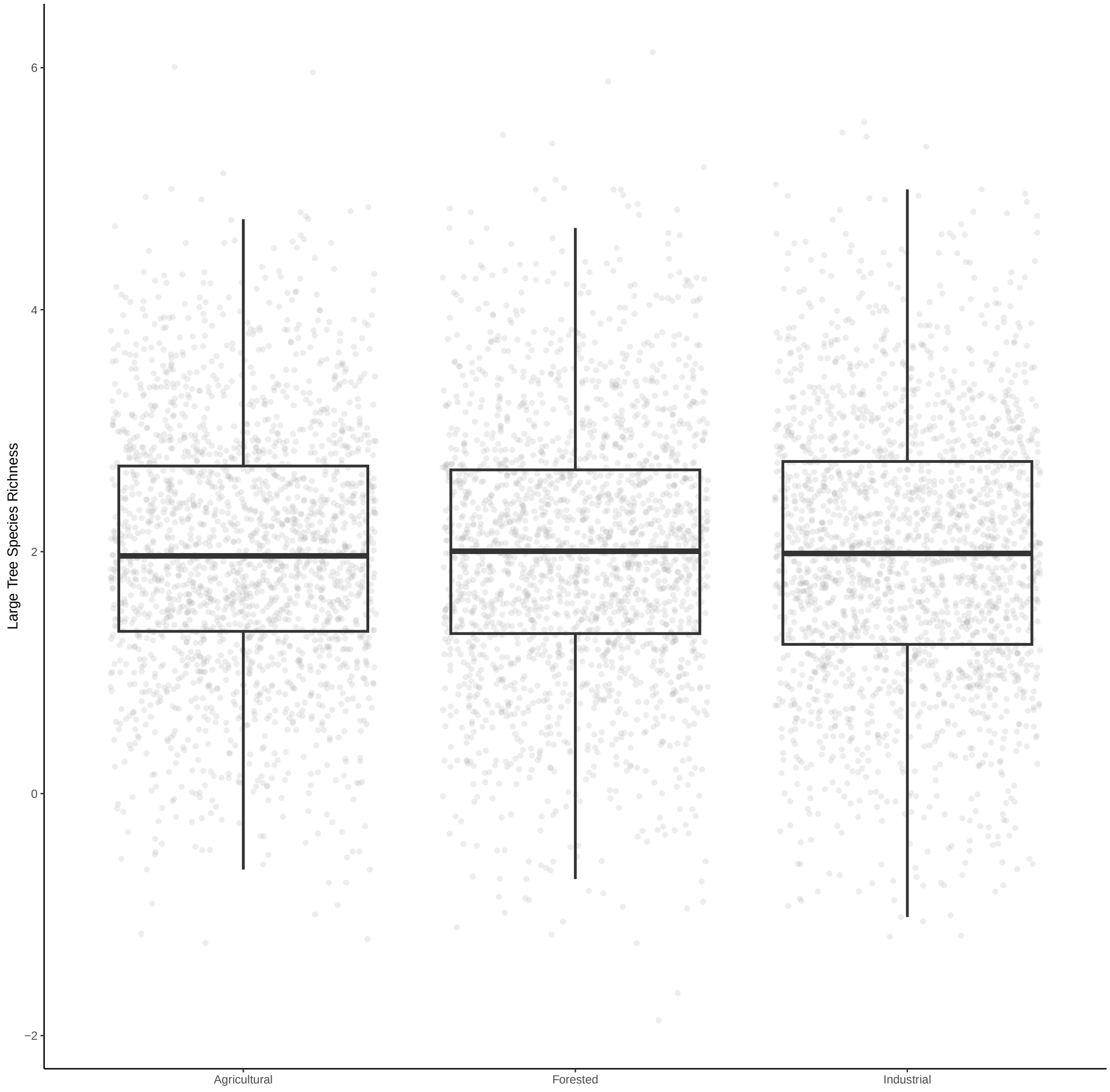

Figure S5f. Model 4 - Direct Effect of Past Land-Use on Large Tree (>= 5 cm DBH) Species Richness

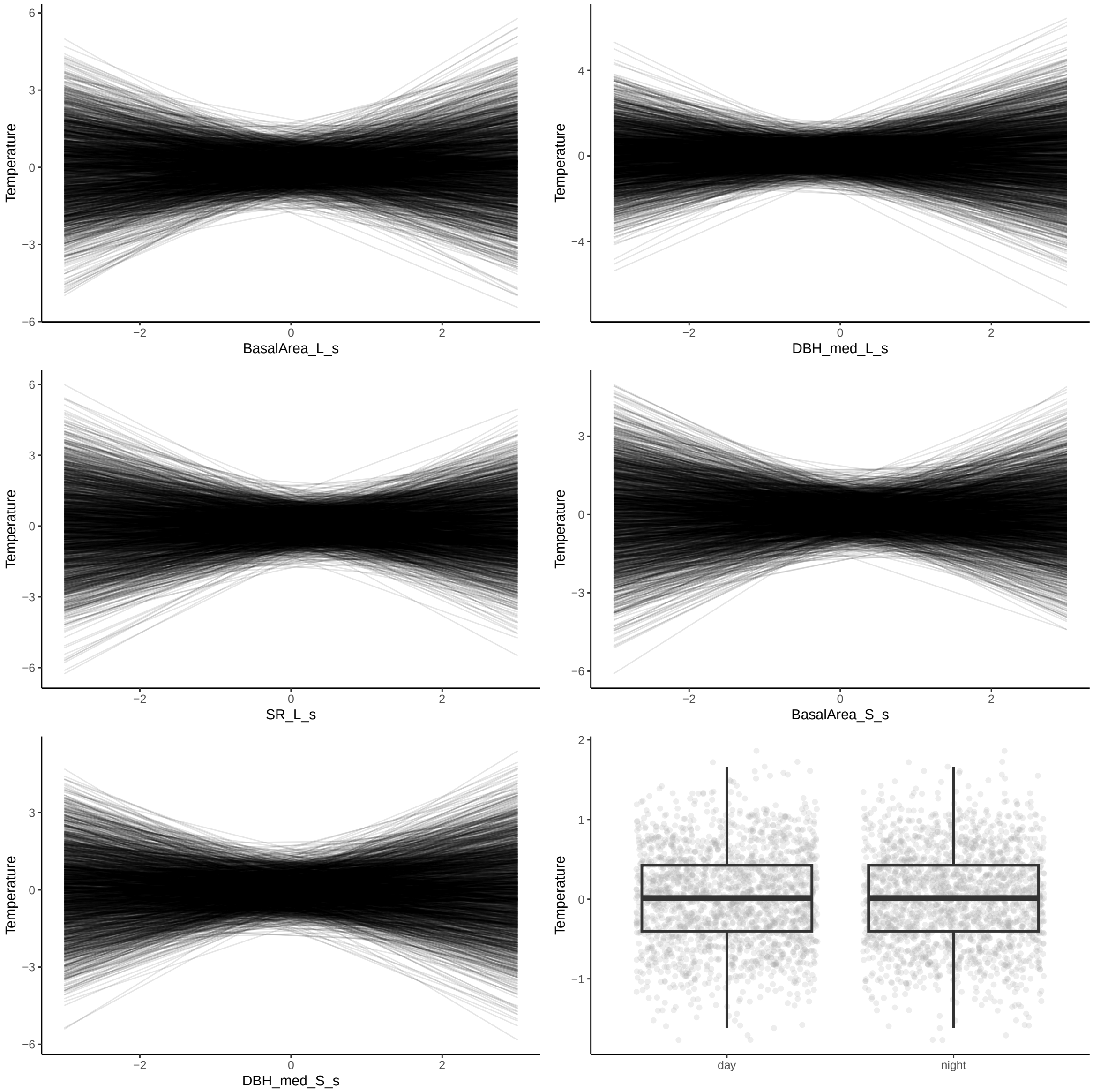

Figure S5g. Model 5 - Direct Effect of Forest Composition on Temperature

### Figure S6

#### Model Diagnostic Plots

These model diagnostic plots assess whether the chains of our models are converged and well mixed, and if the model is well specified and has an adequate fit.

The first plot of the series shows trace plots for each of our parameters, where we want to see stationary and well-mixed chains. The second plot shows an autocorrelation plot by chain and parameter. We want our autocorrelation to quickly drop to zero with increasing lag. Thirdly, the Rhat plot monitors whether a chain has converged to the equilibrium distribution, if all chains are at equilibrium Rhat will be one. If chains have not converged, Rhat will be greater than 1. The fourth plot is the ratio between effective sample size (Neff) and total sample size (N).Because the draws within a Markov chain are not independent if there is autocorrelation, the effective sample size, neff, is usually smaller than the total sample size, N. The larger the ratio, the better. Finally, we have the posterior predictive check where we want the black line to be within/close to the blue lines, to indicate that our model is adequately generative.

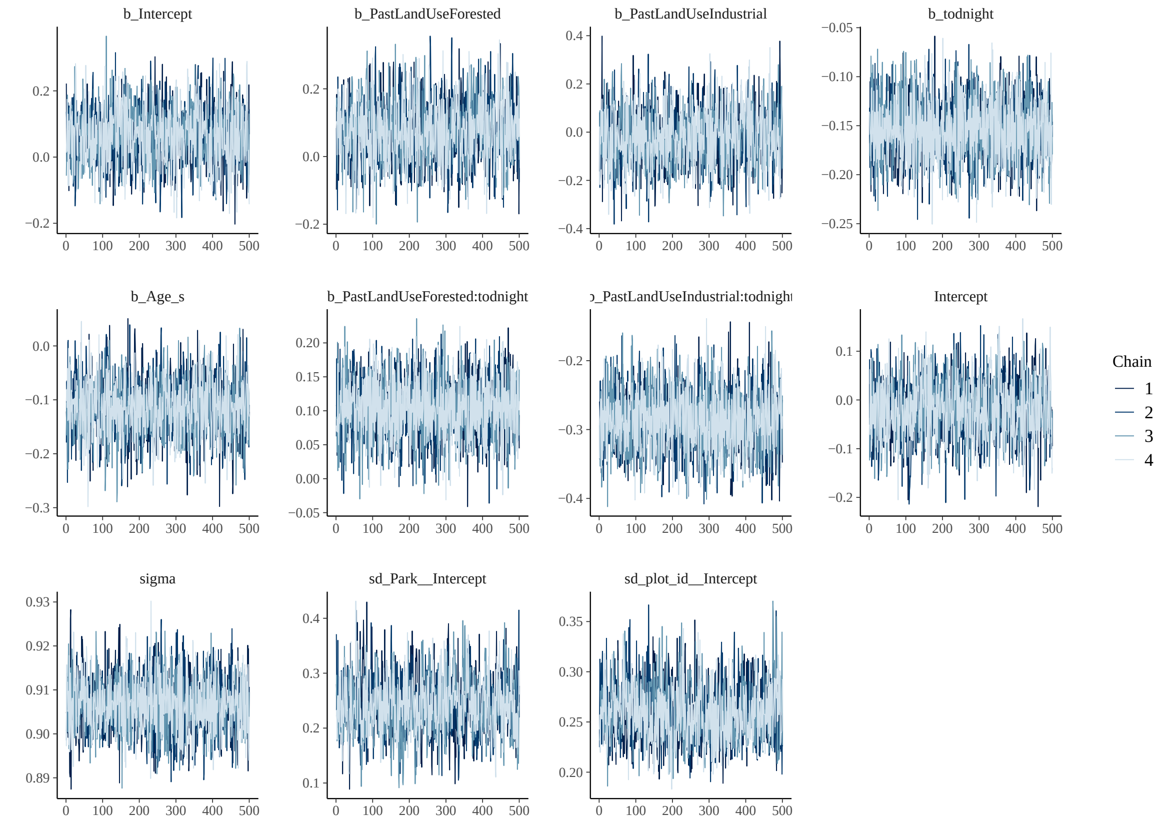
b_Intercept
b_PastLandUseForested
b_PastLandUseIndustrial
b_todnight
b_Age_s
b_PastLandUseForested:todnight
b_PastLandUseIndustrial:todnight
Intercept
sigma
sd_Park__Intercept
sd_plot_id__Intercept
Chain
1
2
3
4

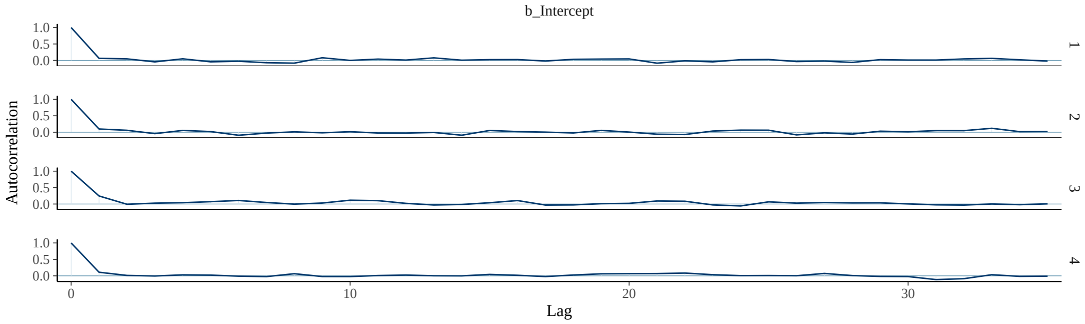
b_Intercept
Autocorrelation
Lag
1
2
3
4

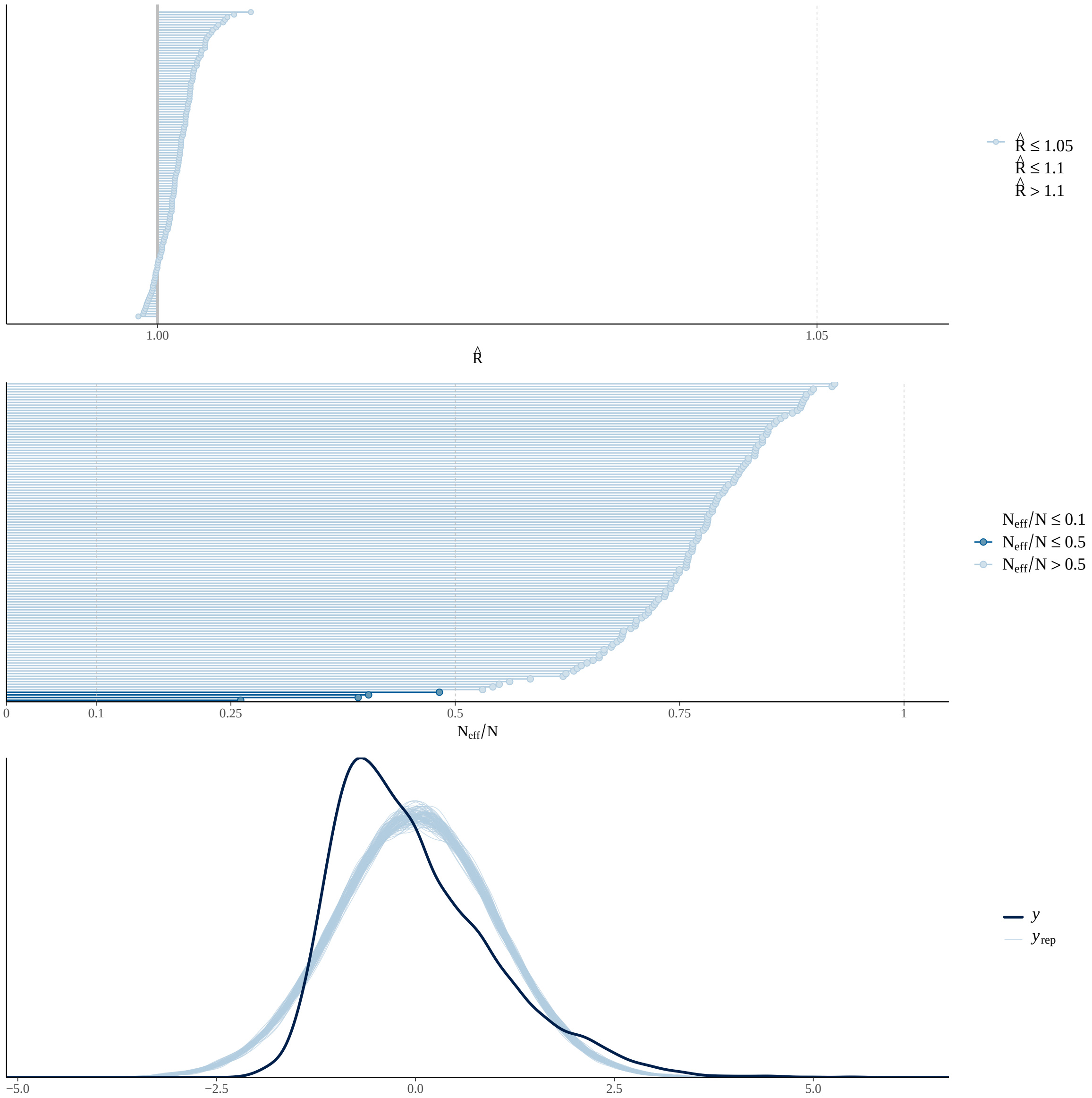

Figure S6a. Model 1 - Total Effect of Past Land-Use on Temperature

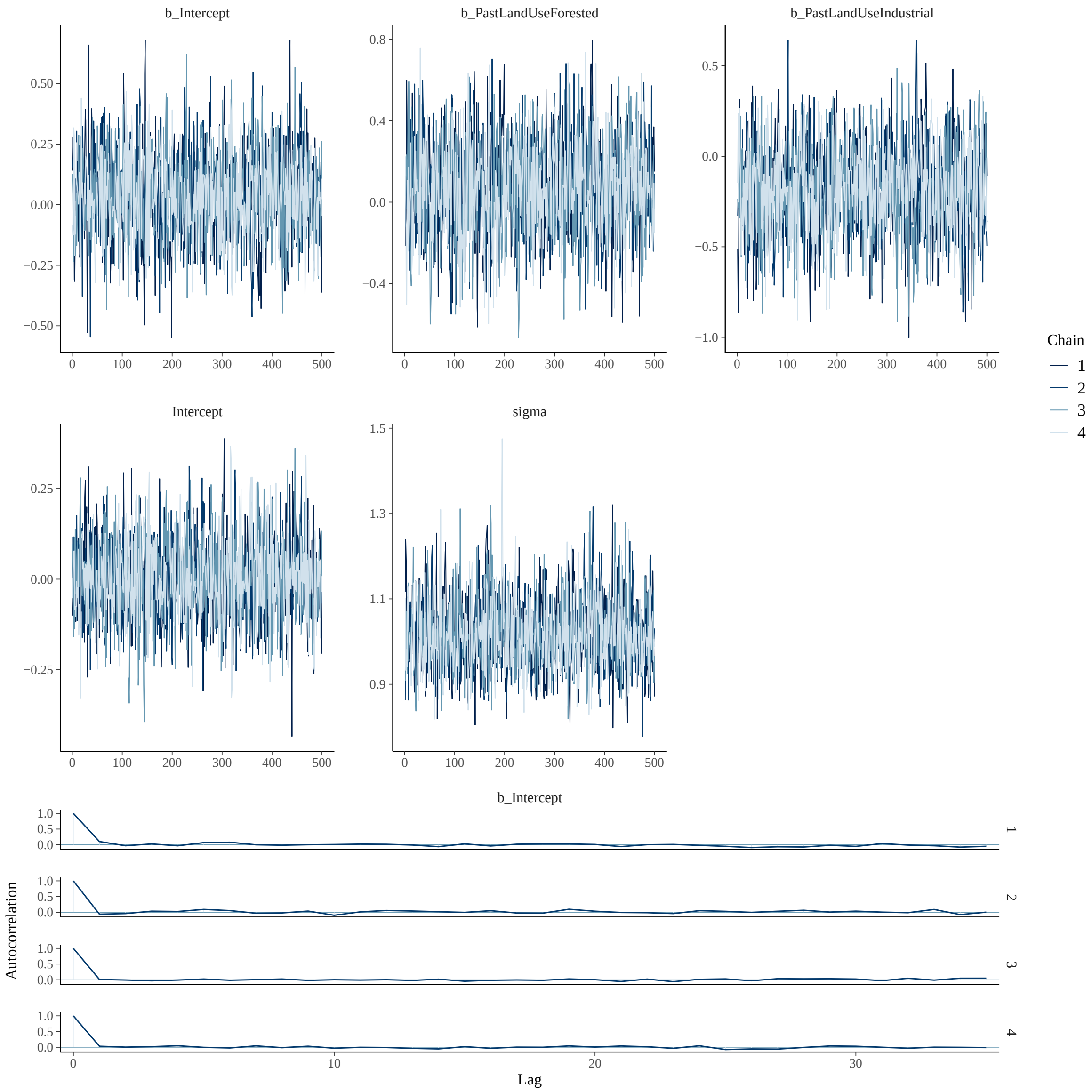

b_Intercept
b_PastLandUseForested
b_PastLandUseIndustrial
Intercept
sigma
Chain
1
2
3
4
b_Intercept
Autocorrelation
Lag

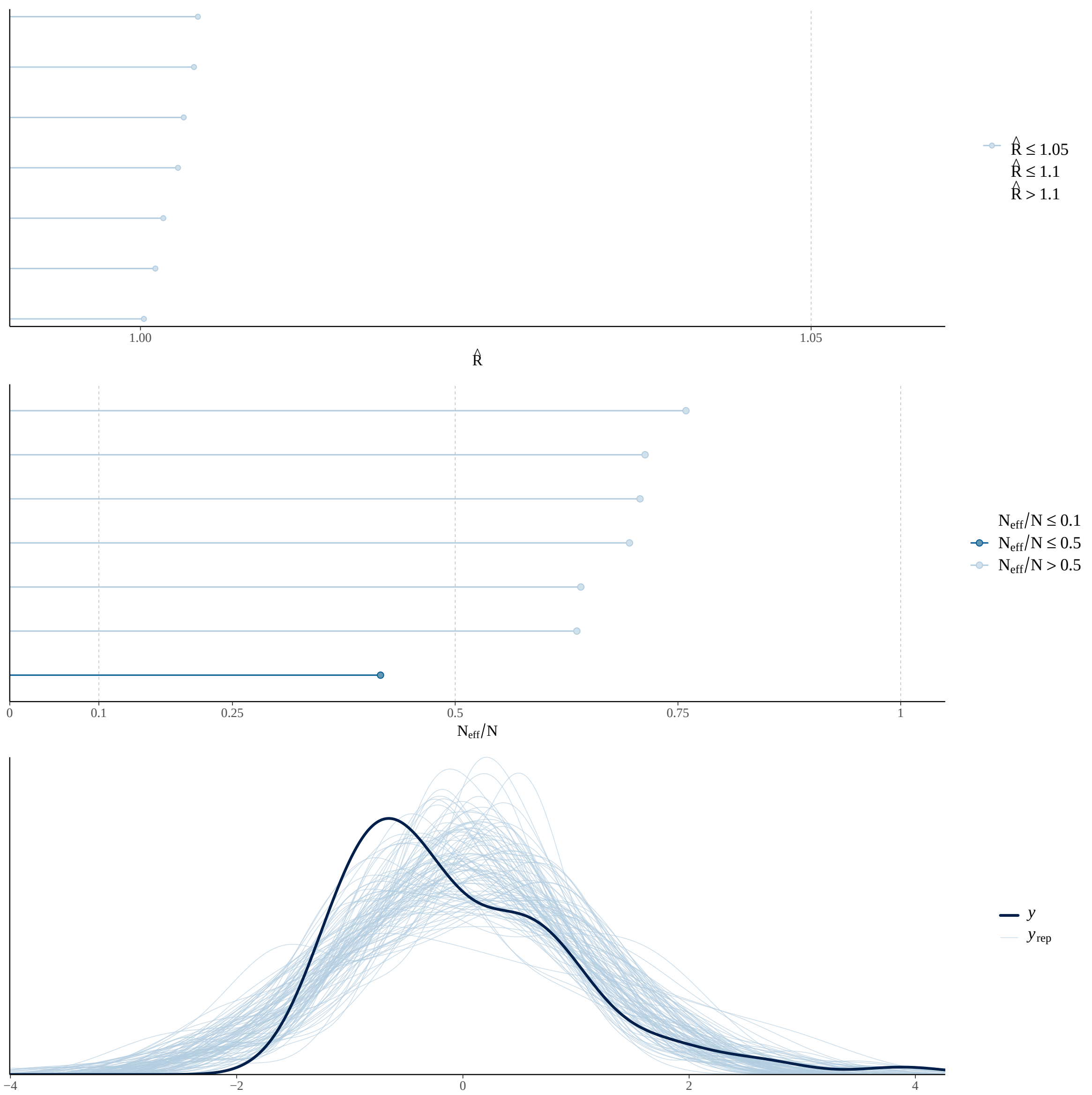

Figure S6b. Model 2a - Direct Effect of Past Land-Use on Large Tree (>= 5 cm DBH) Density

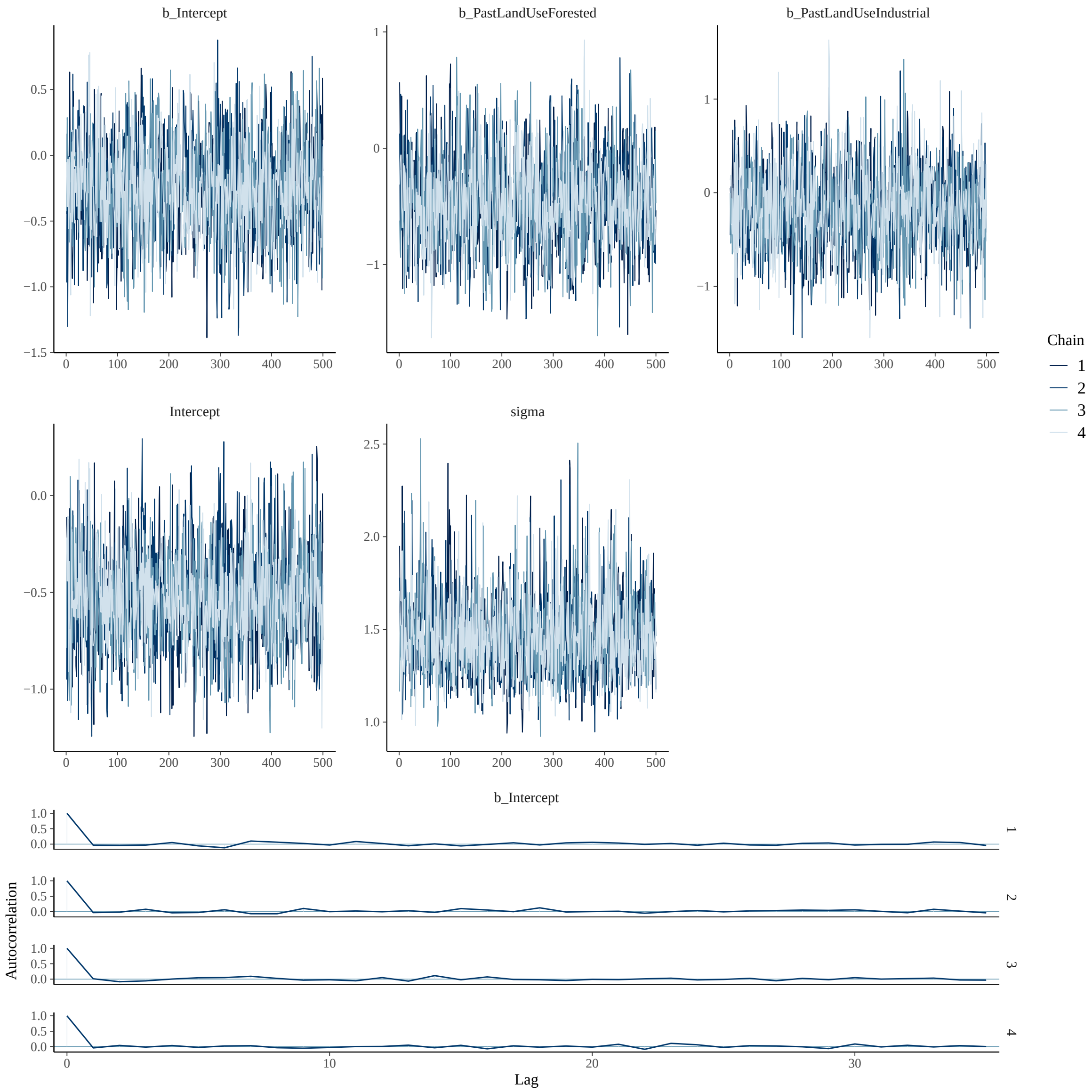
b_Intercept
b_PastLandUseForested
b_PastLandUseIndustrial
Intercept
sigma
Chain
1
2
3
4
b_Intercept
Autocorrelation
Lag

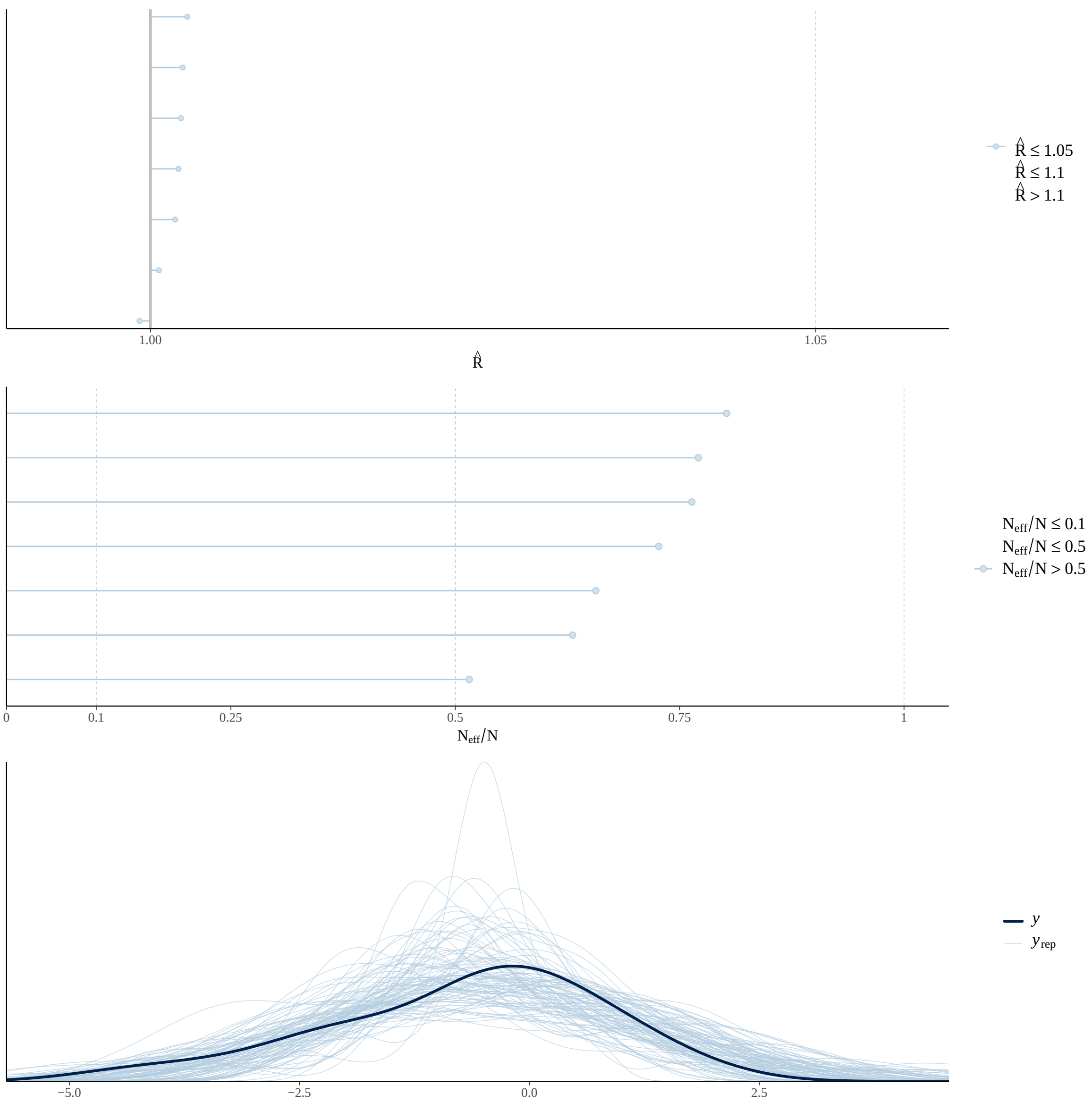

Figure S6c. Model 2b - Direct Effect of Past Land-Use on Small Tree (< 5 cm DBH) Density

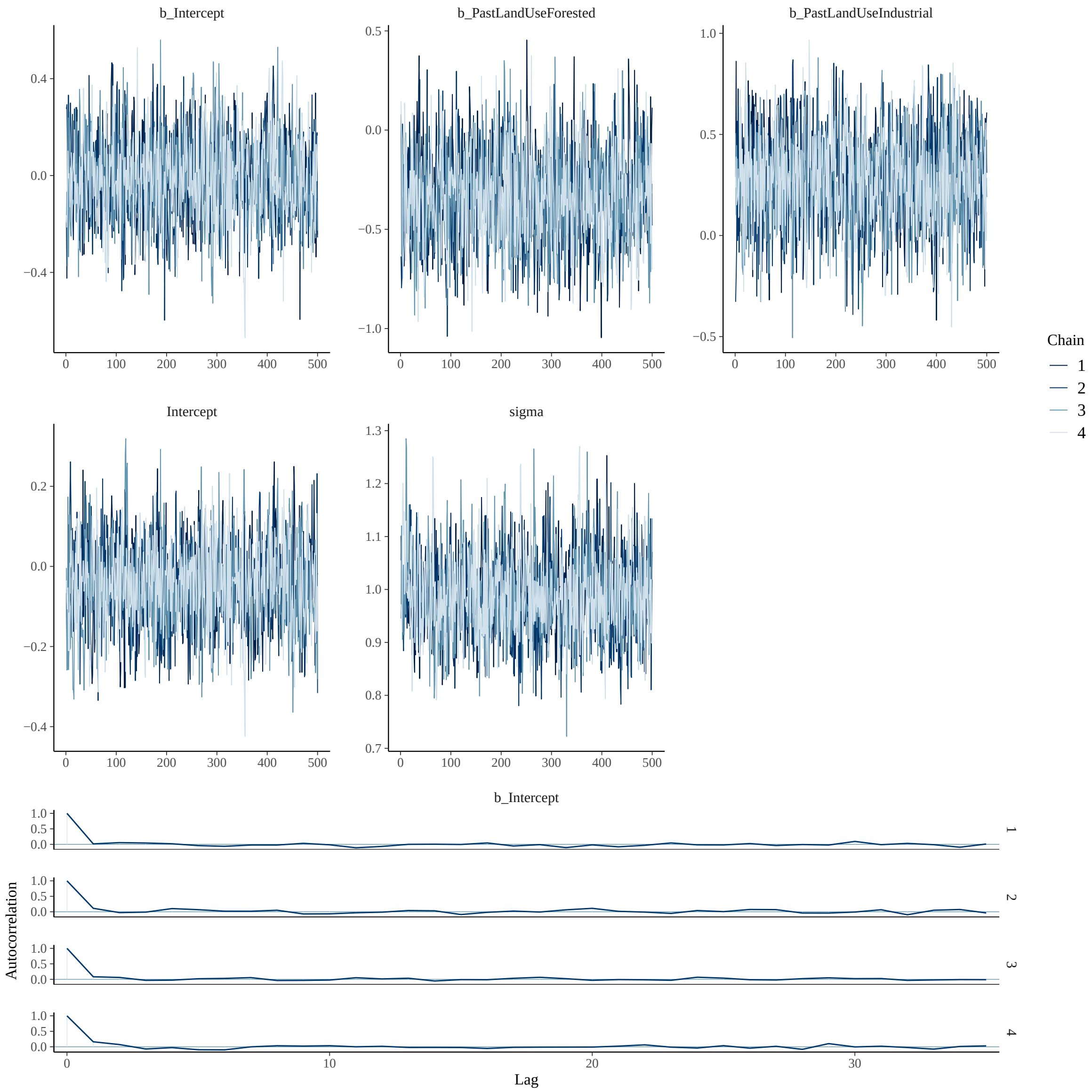
b_Intercept
b_PastLandUseForested
b_PastLandUseIndustrial
Intercept
sigma
Chain
1
2
3
4
b_Intercept
Autocorrelation
Lag

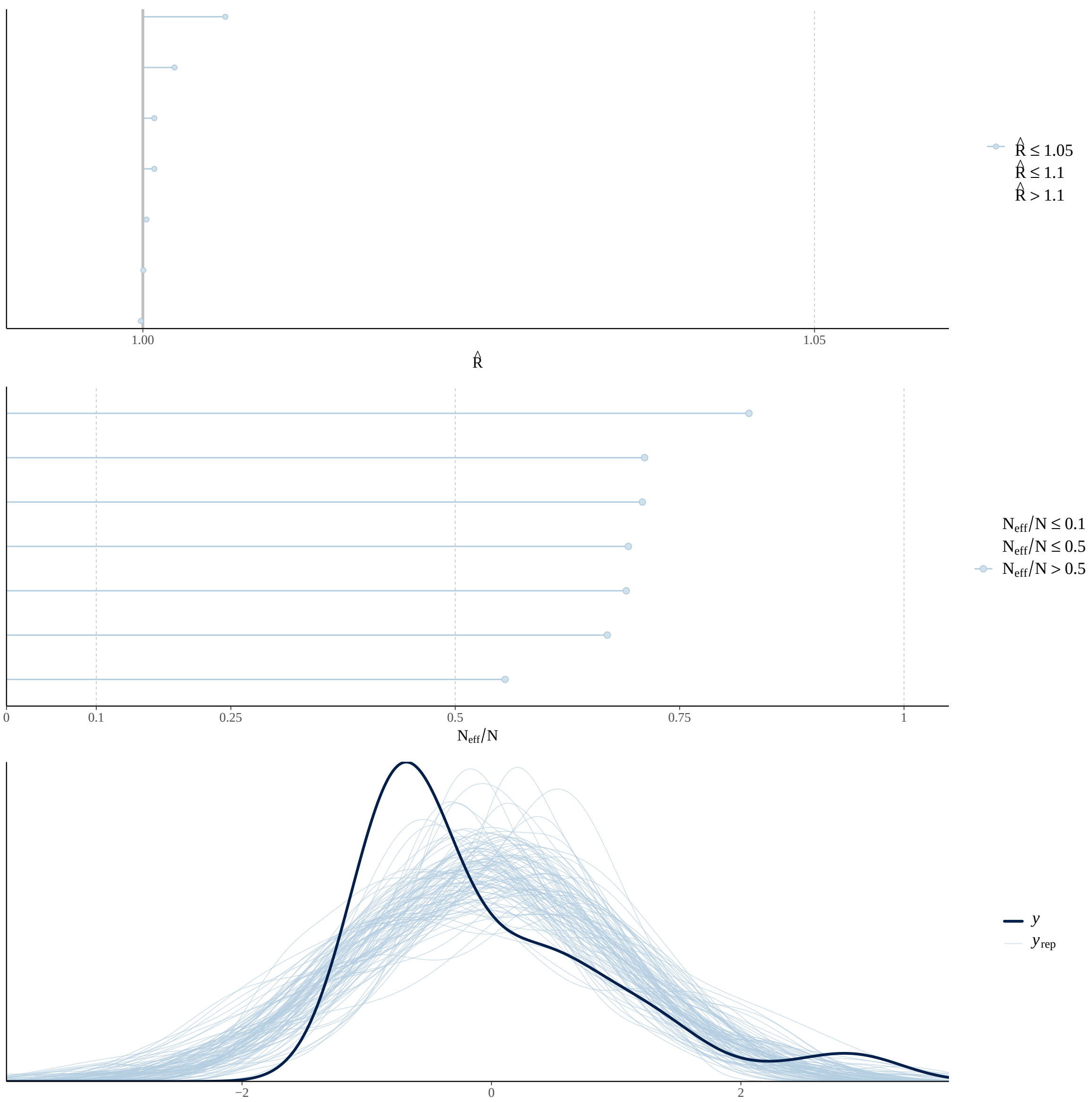

Figure S6d. Model 3a - Direct Effect of Past Land-Use on Large Tree (>= 5 cm DBH) Size

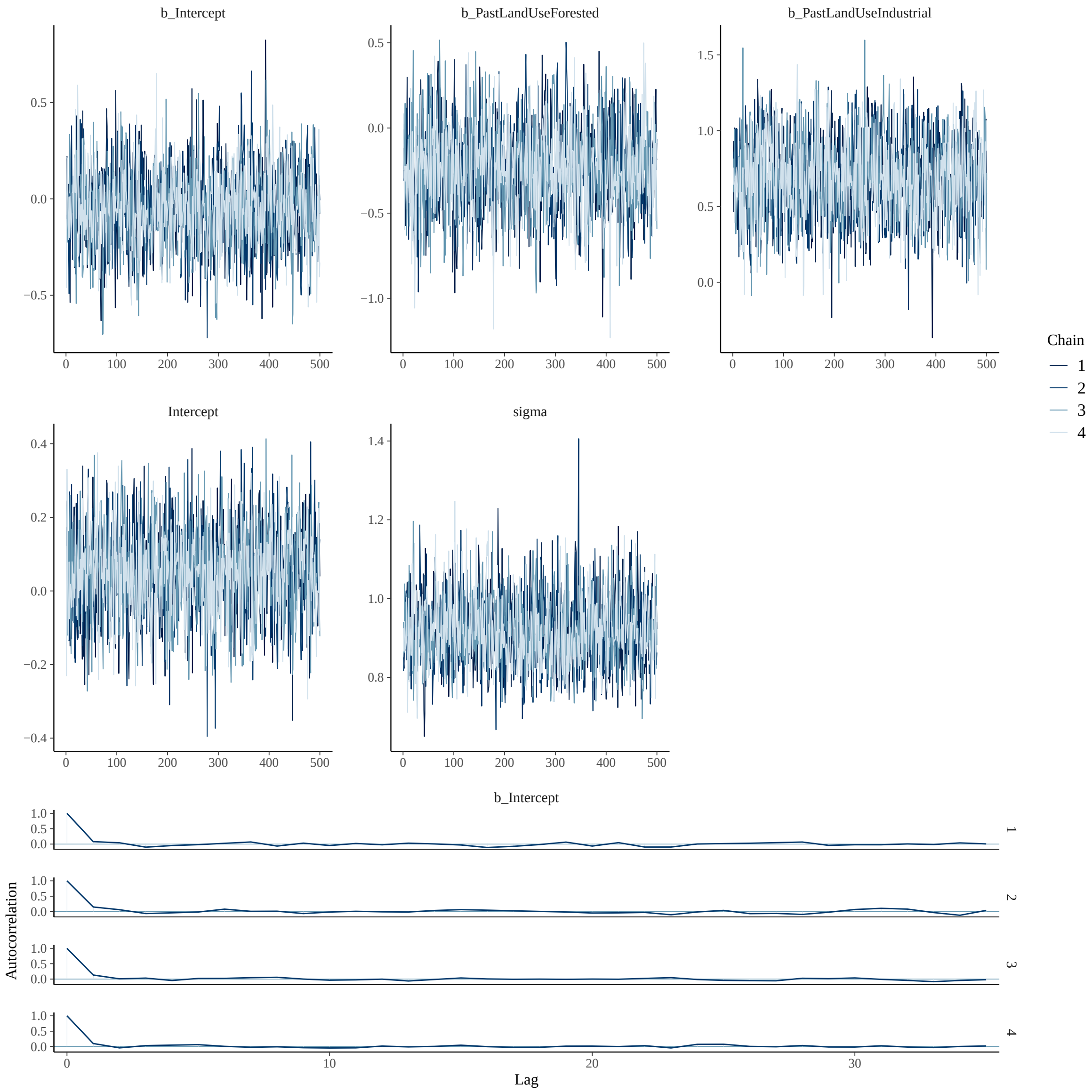

b_Intercept
b_PastLandUseForested
b_PastLandUseIndustrial
Intercept
sigma
Chain
1
2
3
4
b_Intercept
Autocorrelation
Lag

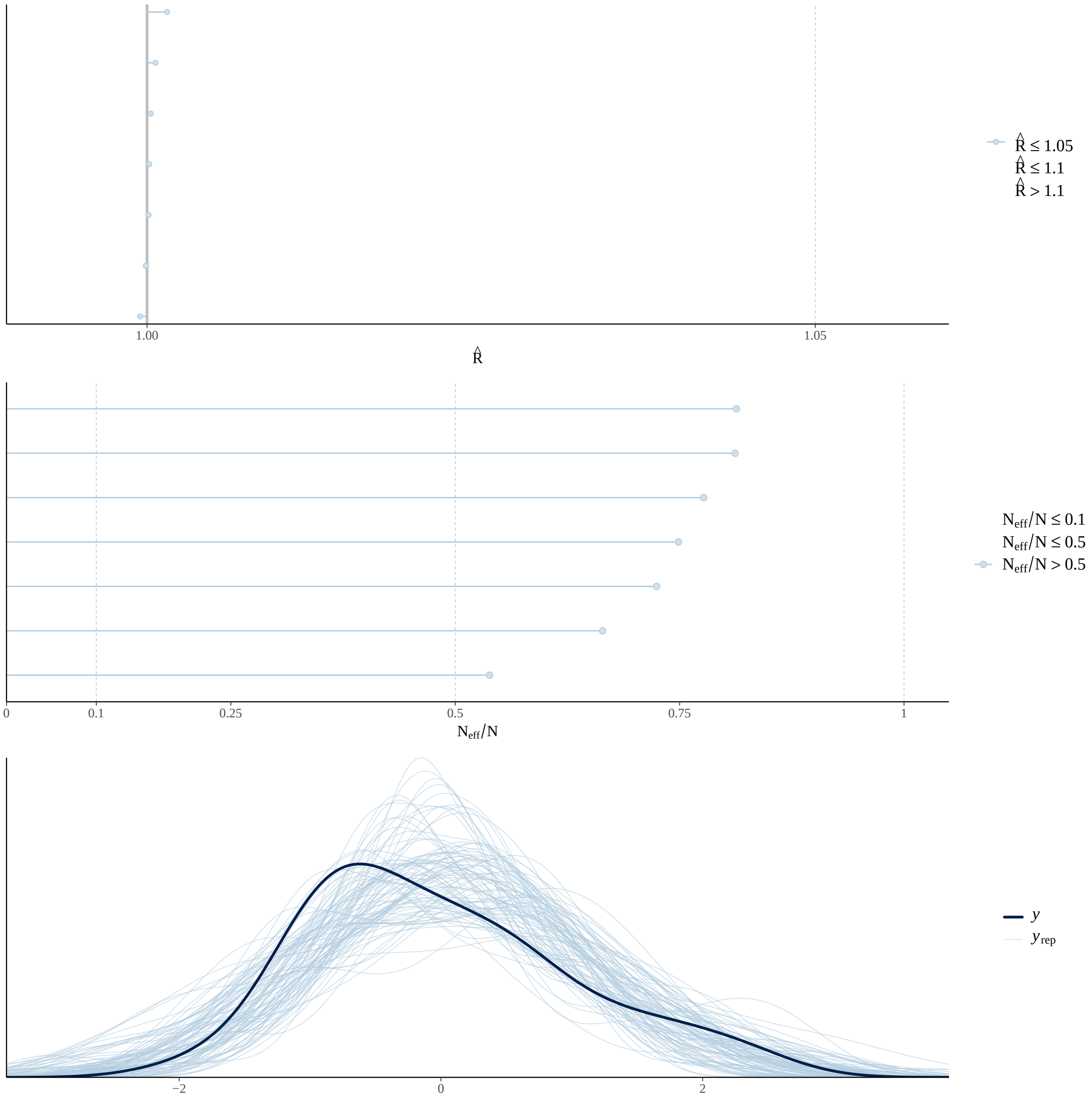

Figure S6e. Model 3b - Direct Effect of Past Land-Use on Small Tree (< 5 cm DBH) Size

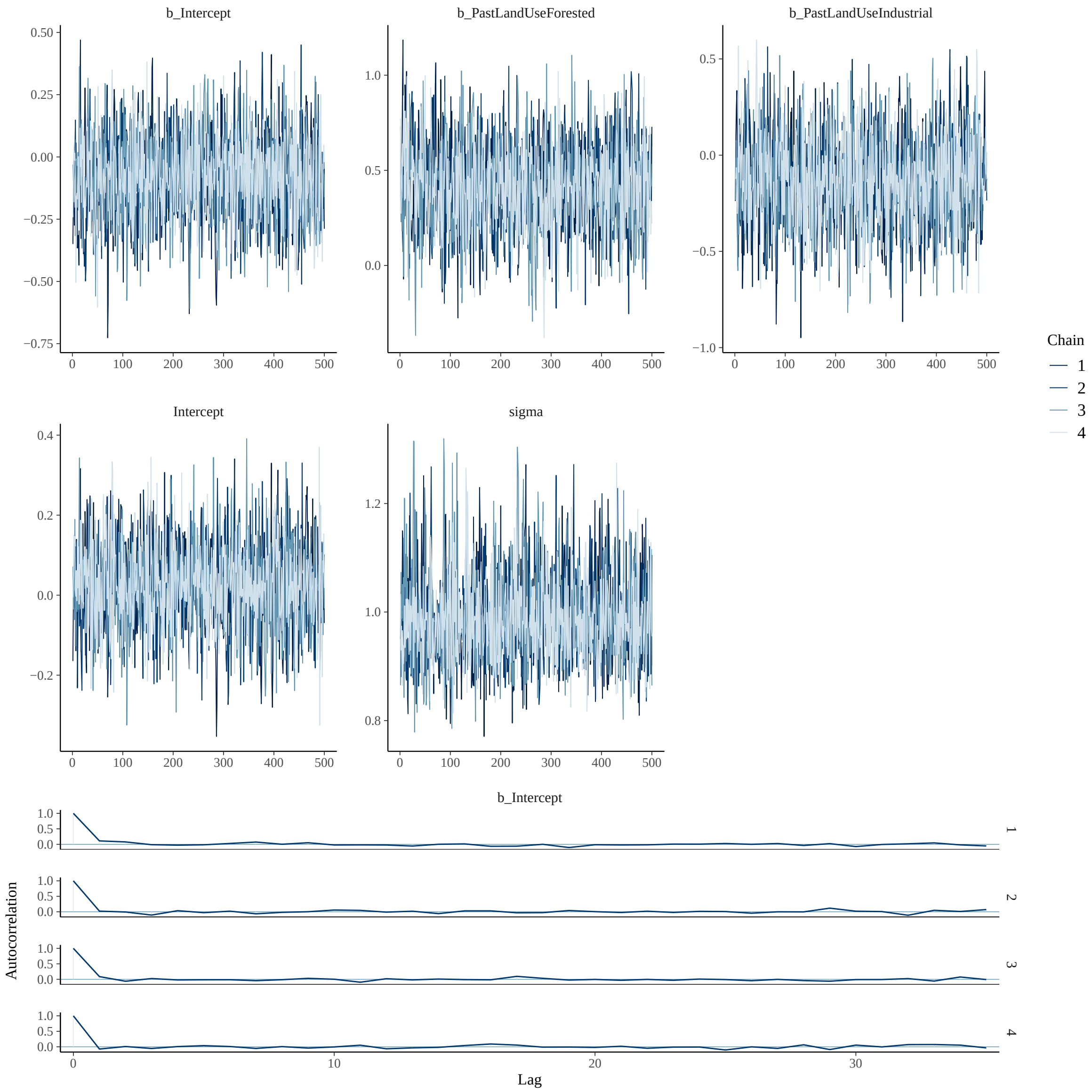
b_Intercept
b_PastLandUseForested
b_PastLandUseIndustrial
Intercept
sigma
Chain
1
2
3
4
Autocorrelation
Lag

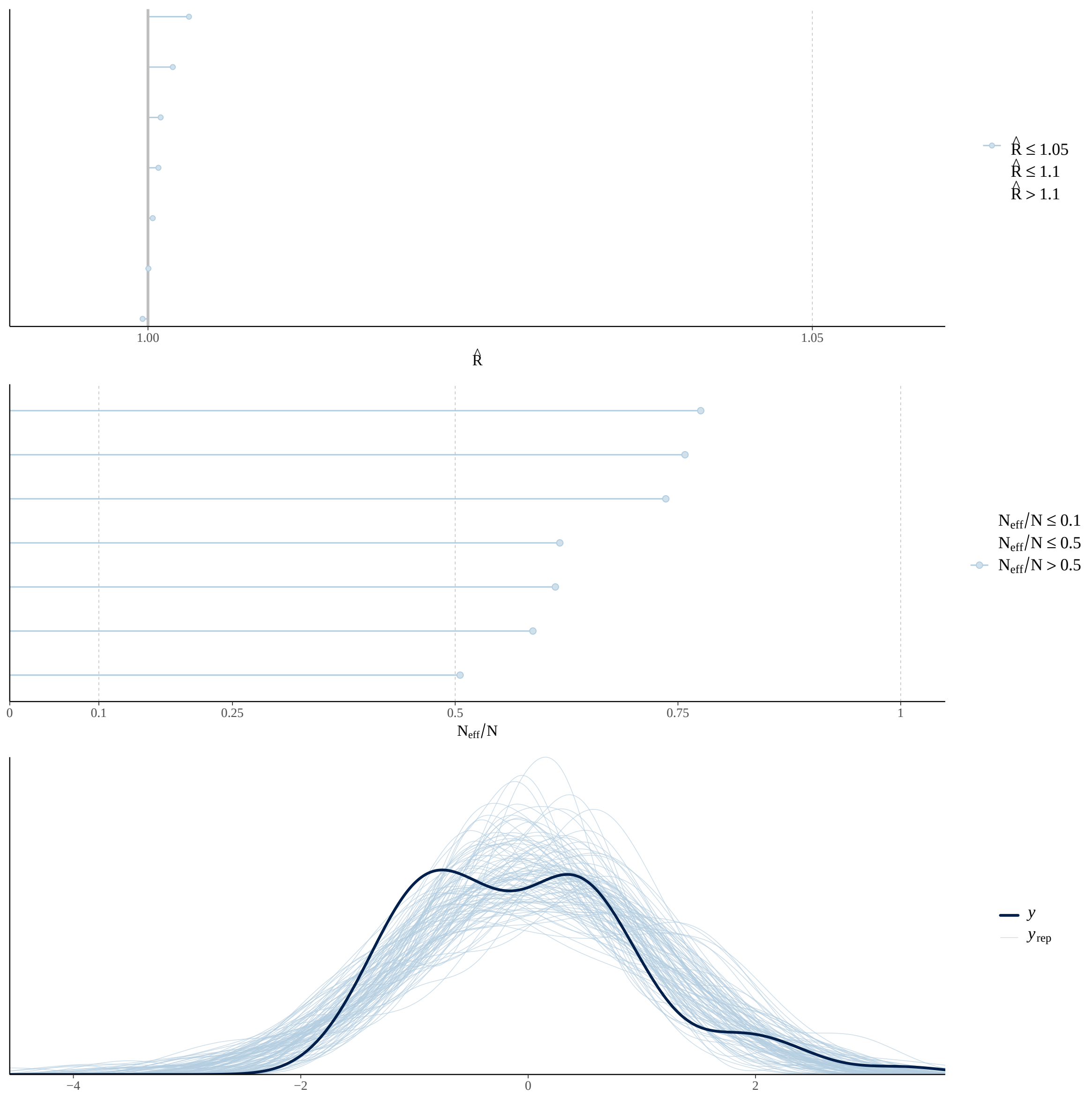

Figure S6f. Model 4 - Direct Effect of Past Land-Use on Large Tree (>= 5 cm DBH) Species Richness

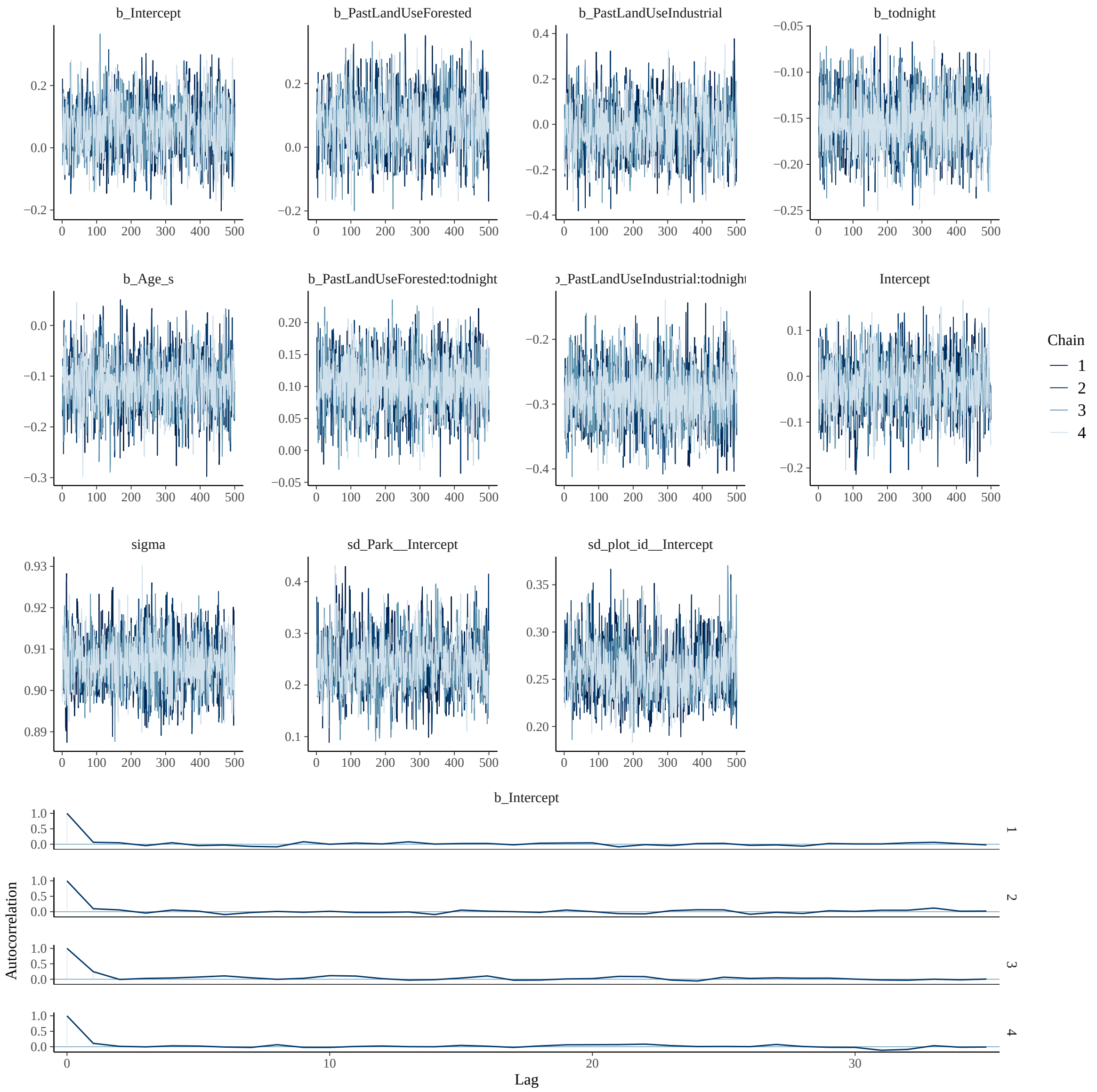
b_Intercept
b_PastLandUseForested
b_PastLandUseIndustrial
b_todnight
b_Age_s
b_PastLandUseForested:todnight
b_PastLandUseIndustrial:todnight
Intercept
sigma
sd_Park__Intercept
sd_plot_id__Intercept
Chain
1
2
3
4
b_Intercept
Autocorrelation
Lag

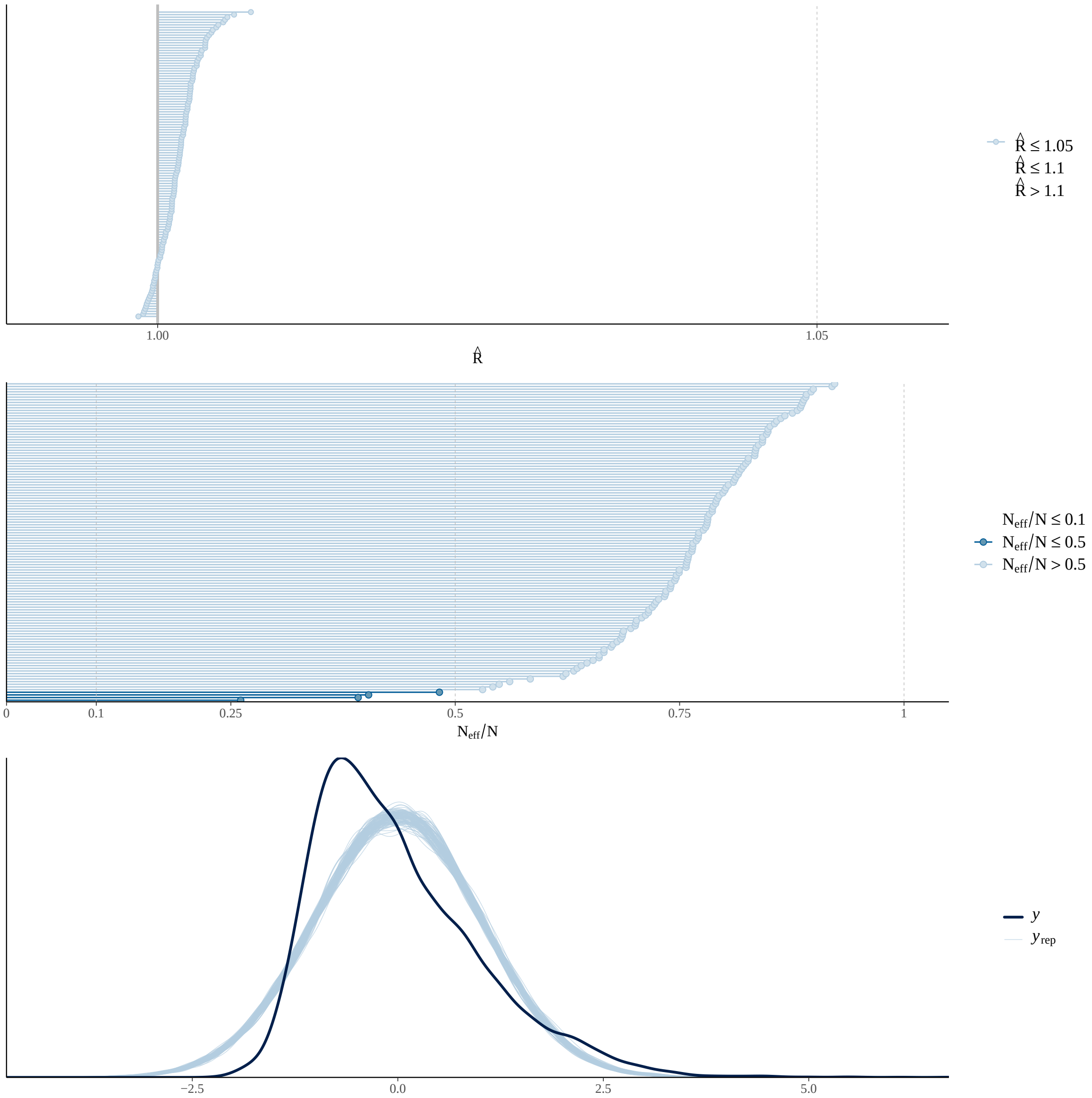

Figure S6g. Model 5 - Direct Effect of Forest Composition on Temperature

### Figure S7

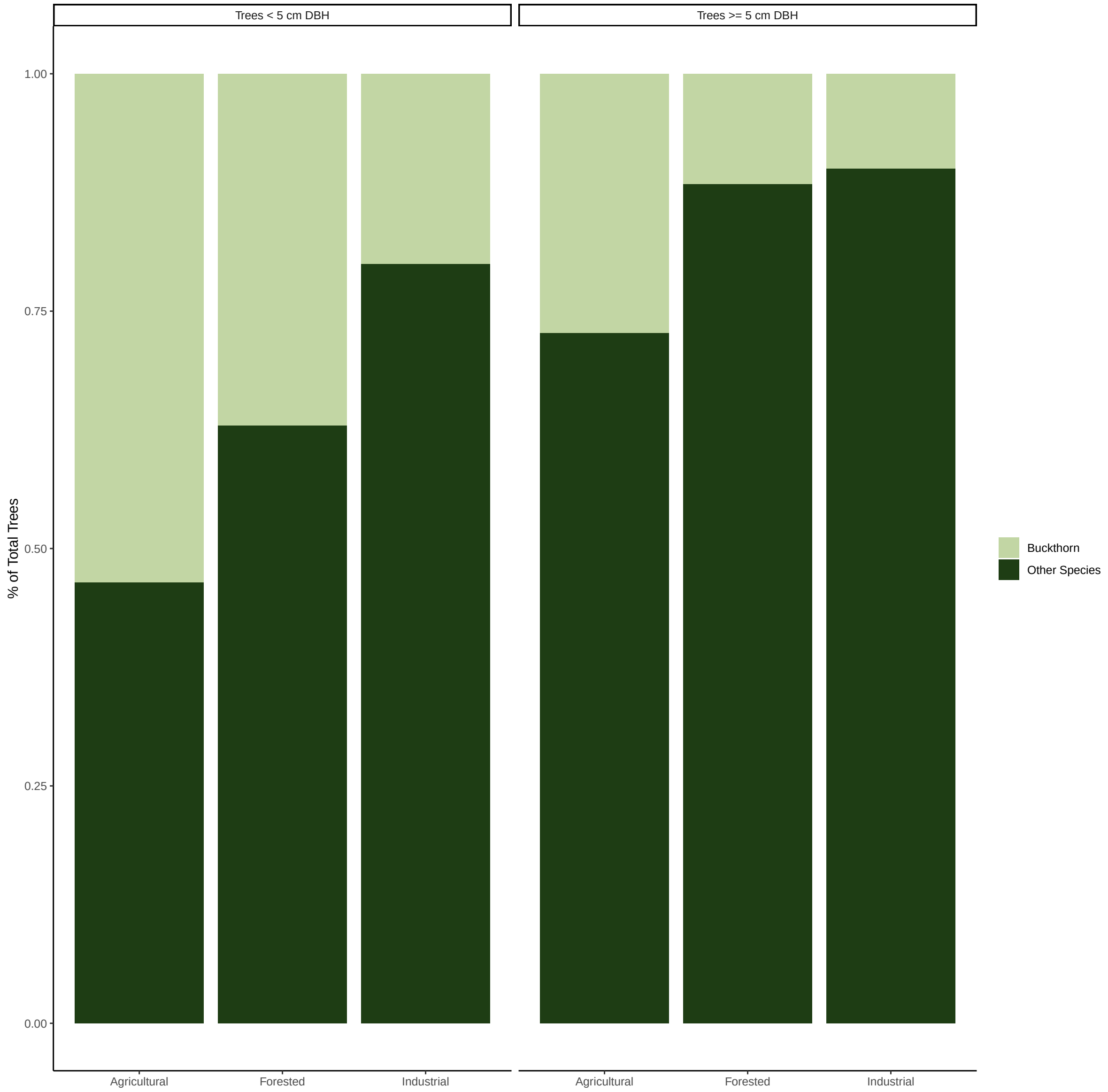

Figure S7. Proportion of common buckthorn (*Rhamnus cathartica*) across past land-use types for trees $<$ 5 cm DBH and trees $>=$ 5 cm DBH. Proportions were calculated across all plots and parks. All other species observed were pooled for the "Other Species" category.

#### Figure S8

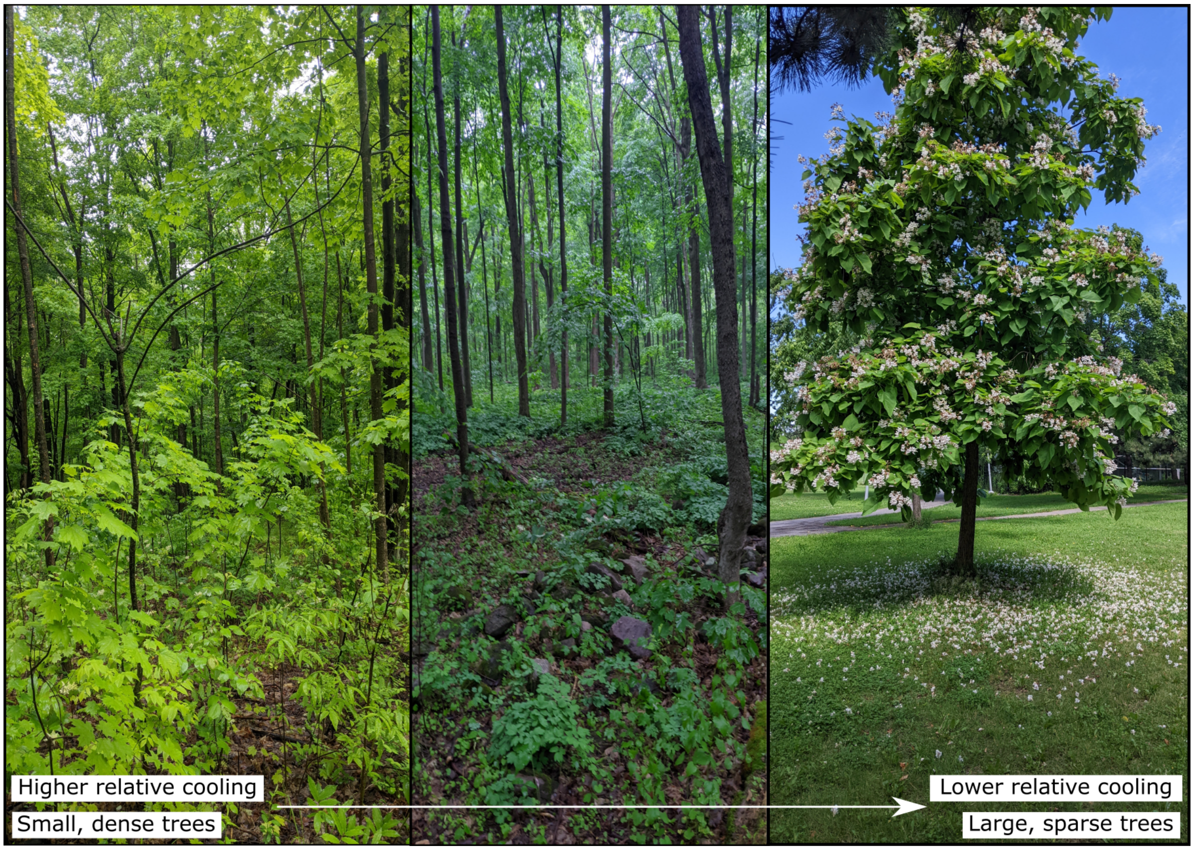

Figure S8. Examples of the trade-off that happens in urban parks, moving from areas with small, dense trees (often in previously forested or agricultural "nature" parks) to large, sparse trees (often found in more traditional "urban" parks). Photos taken by Isabella C Richmond.

---

### Tables

#### Table S1.

Table S1. Historical data sources used to reconstruct history of park sites to extract previous land-cover type and age of park.

| Title | Organization | Data.Type | Date | Link |
| --- | --- | --- | --- | --- |
| D1900-1 : Parcs - Administration | Archives de Montréal | Historical documentation | [18-]-[19-] | https://archivesdemontreal.ica-atom.org/d1900-1-parcs-administration-18-19 |
| D1901-18-1 : Parcs et terrains de jeux : Parc Baldwin - Historique. | Archives de Montréal | Historical documentation | [18-]-[19-] | https://archivesdemontreal.ica-atom.org/d1901-18-1-parcs-et-terrains-de-jeux-parc-baldwin-historique-18-19 |
| D1901-38-30-3 : Parcs et terrains de jeux : Parc La Fontaine - Historique - Documents. | Archives de Montréal | Historical documentation | [18-]-[19-] | https://archivesdemontreal.ica-atom.org/d1901-38-30-3-parcs-et-terrains-de-jeux-parc-la-fontaine-historique-documents-18-19 |
| D1925-2 : Parcs - Jardin botanique : Historique. | Archives de Montréal | Historical documentation | [18-]-[19-] | https://archivesdemontreal.ica-atom.org/d1925-2-parcs-jardin-botanique-historique-19 |
| D1925-3 : Parcs - Jardin botanique : Construction. | Archives de Montréal | Historical documentation | [19-] | https://archivesdemontreal.ica-atom.org/d1925-3-parcs-jardin-botanique-construction-19 |
| D3040-12-A : Ile Bizard : Général. | Archives de Montréal | Historical documentation | [19-] | https://archivesdemontreal.ica-atom.org/d3040-12-a-ile-bizard-general-19 |
| D3050-23-19 : Ville - Pierrefonds : Parcs. | Archives de Montréal | Historical documentation | [19-] | https://archivesdemontreal.ica-atom.org/d3050-23-19-ville-pierrefonds-parcs-19 |

*(continued)*

| Title | Organization | Data.Type | Date | Link |
|---|---|---|---|---|
| D1900-A-4 : Parcs et terrains de jeux - Répertoire alphabétique : Général. | Archives de Montréal | Historical documentation | Après 1957 | https://archivesdemontreal.ica-atom.org/d1900-a-4-parcs-et-terrains-de-jeux-repertoire-alphabetique-general-apres-1957 |
| D1900-A-4 : Parcs et terrains de jeux - Répertoire alphabétique : Général. | Archives de Montréal | Historical documentation | Avant 1958 | https://archivesdemontreal.ica-atom.org/d1900-a-4-parcs-et-terrains-de-jeux-repertoire-alphabetique-general-avant-1958 |
| D1901-17-1 : Parcs et terrains de jeux - Île Sainte-Hélène : Historique. | Archives de Montréal | Historical documentation | Avant 1961 | https://archivesdemontreal.ica-atom.org/d1901-17-1-parcs-et-terrains-de-jeux-ile-sainte-helene-historique-avant-1961 |
| D1900-A-1 : Parcs et terrains de jeux : Général. | Archives de Montréal | Historical documentation | Après 1971 | https://archivesdemontreal.ica-atom.org/d1900-a-4-parcs-et-terrains-de-jeux-repertoire-alphabetique-general-avant-1958 |
| D1900-A-1 : Parcs et terrains de jeux : Général. | Archives de Montréal | Historical documentation | Avant 1972 | https://archivesdemontreal.ica-atom.org/d1900-a-1-parcs-et-terrains-de-jeux-general-avant-1972 |
| Photothèque 1930-2003 | Division de la géomatique | Satellite imagery | 1930-2003 | https://www.arcgis.com/apps/webappviewer/in[illegible] |

*(continued)*

| Title | Organization | Data.Type | Date | Link |
|---|---|---|---|---|
| Carte de localisation des anciennes carrières et des dépôts de surface de la Ville de Montréal | Division Soutien Technique, Infrastructures, CESM | Historical documentation, spatial data | 2023 | https://www.arcgis.com/apps/webappviewer/in |
| Angrignon | Personal website | Historical documentation | | https://www.angrignon.ca/gene/indexg1.html |
| Analyse de la valeur patrimoniale de la Maison Jacques Richer dit Louveteau | Arrondissement Pierrefonds-Roxboro | Historical documentation | 2008 | http://ville.montreal.qc.ca/pls/portal/docs/PA |
| Le parc Thomas-Chapais | WWF | Historical documentation | 2016 | https://wwf.ca/wp-content/uploads/2016/11/Thomas-Chapais_Caract%C3%A9ristiques_%C3%A9co |
| Historique de Tétreaultvill | Comité citoyen du parc Thomas-Chapais | Historical documentation | | https://comitecitoyenduparcthomas-chapais.ca/historique/ |
| Le Douglas: une longue histoire d'excellence | Douglas Research Centre | Historical documentation | | https://douglas.research.mcgill.ca/fr/historique |
| Le boisé de Saint Sulpice | Blog | Historical documentation | 2020 | https://floraurbana2.blogspot.com/2020/06/le-boise-de-saint-sulpice.html |

*(continued)*

| Title | Organization | Data.Type | Date | Link |
|---|---|---|---|---|
| Ancien Dépotoir du Boisé de l'Heritage | Gestion des Lieux Contaminés | Historical documentation | 1987 | https://drive.google.com/file/u/1/d/133JOw0G |
| Ancien Dépotoir Beaubien | Gestion des Lieux Contaminés | Historical documentation | 1991 | https://drive.google.com/file/u/1/d/133JOw0G |
| Extrémité Sud et Sud-Ouest de l'Ile-des-Soeurs | Gestion des Lieux Contaminés | Historical documentation | 1991 | https://drive.google.com/file/u/1/d/133JOw0G |
| Commission de toponymie | Gouvernement du Québec | Historical documentation | 2023 | https://toponymie.gouv.qc.ca/ct/ |
| Maison Saint-Gabriel | Grand répertoire du patrimoine bati de Montréal | Historical documentation | 2012 | http://patrimoine.ville.montreal.qc.ca/inventair |
| Ancien Dépotoir du Parc Lafond | Group d'étude et de restauration des lieux d'élimination des déchets dangereux | Historical documentation | | https://drive.google.com/file/d/1egb2BQx4WX BPaGbY/view |
| Détection de biogaz au Parc Baldwin | Ville de Montréal | Historical map | 1992 | https://drive.google.com/file/u/1/d/1HkSGrBr |

*(continued)*

| Title | Organization | Data.Type | Date | Link |
|---|---|---|---|---|
| Parc-nature de l'Anse-à-l'Orme | Laboratoire d'histoire et de patrimoine de Montréal | Historical documentation | | https://chronomontreal.uqam.ca/chronologie/2[illegible] parc-nature-de-l-anse-a-l-orme |
| Parc-nature du Bois-de-Liesse | Laboratoire d'histoire et de patrimoine de Montréal | Historical documentation | | https://montreal.ca/toponymie/toponymes/par[illegible] regional-du-bois-de-liesse |
| History of Centennial Hall and | | | | |
| Memorial Park | Personal report | Historical documentation | 2013 | https://www.shbbhs.ca/j3/images/RAngellCon[illegible] |
| Fritz Farm Planning Committee, N&V, Sep 1980 | Town Hall report | Historical documentation | 1980 | http://home.linuxfocus.org/~guido/book-a-history-of-the-town-of-baie-d-urfe/then-and-now/2020-08-09_1734-088-nv-sep-1980-fritz-park-planing.html |
| Le Domaine Saint-Sulpice, un quartier en partage | Société D'Histoire d'Ahuntsic-Cartierville | Historical documentation | 2015 | https://www.lashac.com/panneaux-historiques.html |
| Adrien-D. Archambault | Société Historique de Montréal | Historical documentation | 2020 | https://www.societehistoriquedemontreal.org/n[illegible] histoire/liste-des-presidents-depuis-la-fondation/adrien-d-archambault/ |

*(continued)*

| Title | Organization | Data.Type | Date | Link |
|---|---|---|---|---|
| Parcours riverain | Ville de Montréal | Historical documentation, spatial data | 2023 | https://donnees.montreal.ca/dataset/parcours-riverain |
| Répertoire historique des toponymes | Ville de Montréal | Historical documentation | 2023 | https://montreal.ca/toponymie/ |
| Le parc Frédéric-Back : une métamorphose unique | Ville de Montréal | Historical documentation | 2023 | https://montreal.ca/articles/le-parc-frederic-back-une-metamorphose-unique-18997 |

### Table S2

Table S2. Park names, ID, legacies (i.e., past land use(s)), the municipalities parks are found in, and if they are in the City of Montreal, their borough, the year they were established, the number of temperature sensors deployed in each park, and the number of temperature sensors retrieved at the end of the season. An ID that begins with CON indicates a control sensor, found in a parking lot at least 500 m away from the associated park(s). Benny park had all three sensors stolen and was thus removed from the temperature analysis.

| Park | Park.ID | Legacy.ies. | Municipality | Borough | Year Established | Sensors Deployed | Sensors Retrieved |
|---|---|---|---|---|---|---|---|
| Adrien-D-Archambault | IND1 | Industrial | Montreal | Verdun | 2002 | 3 | 3 |
| Angrignon | FOR1, AGR1 | Forested, Agricultural | Montreal | Le Sud-Ouest | 1927 | 6 | 5 |
| Arborétum du Jardin botanique | IND7 | Industrial | Montreal | Rosemont-La Petite-Patrie | 1936 | 3 | 3 |
| Arthur-Therrien | IND2 | Industrial | Montreal | Verdun | ~ 1964 | 3 | 3 |

*(continued)*

| Park | Park.ID | Legacy.ies. | Municipality | Borough | Year Established | Sensors Deployed | Sensors Retrieved |
|---|---|---|---|---|---|---|---|
| Baldwin | IND3 | Industrial | Montreal | Le Plateau-Mont Royal | 1909 | 3 | 3 |
| Benny | AGR7 | Agricultural | Montreal | Notre-Dame-de-Grâce | 1965 | 3 | 0 |
| Boisé-du-Saint-Sulpice | FOR4 | Forested | Montreal | Ahuntsic-Cartierville | 1990 | 3 | 3 |
| Bois-de-Liesse | FOR2 | Forested | Montreal | Pierrefonds-Roxboro / Saint-Laurent | 1987 | 3 | 3 |
| Bois-de-Saraguay | FOR3 | Forested | Montreal | Ahuntsic-Cartierville | 1987 | 3 | 3 |
| Cap-Saint-Jacques | FOR5, AGR2 | Forested, Agricultural | Montreal | Pierrefonds-Roxboro | 1985 | 6 | 6 |
| Centennial Hall | AGR8 | Agricultural | Beaconsfield | | 1945 | 3 | 3 |
| Coulée-Grou | FOR6 | Forested | Montreal | Rivière-des-Prairie-Pointe-aux-Trembles | 1988 | 3 | 3 |
| Félix-Leclerc | IND5 | Industrial | Montreal | Mercier-Hochelaga-Maisonneuve | 1990 | 3 | 2 |
| Frédéric-Back | IND6 | Industrial | Montreal | Vileray-Saint Michel-Parc Extension | 2016 | 3 | 2 |
| Fritz | AGR9 | Agricultural | Baie-d’Urfe | | 1979 | 3 | 3 |
| Île-Bizard | FOR7, AGR3 | Forested, Agricultural | Montreal | L’Île-Bizard–Sainte-Geneviève | 1991 | 6 | 6 |

*(continued)*

| Park | Park.ID | Legacy.ies. | Municipality | Borough | Year Established | Sensors Deployed | Sensors Retrieved |
|---|---|---|---|---|---|---|---|
| Jean-Drapeau | FOR8 | Forested | Montreal | Ville-Marie | 1908 | 3 | 3 |
| L'Anse-à-L'Orme | FOR9, AGR4 | Forested, Agricultural | Montreal | Pierrefonds-Roxboro / Saint-Laurent | 1987 | 6 | 6 |
| Lafond | IND8 | Industrial | Montreal | Rosemont-La Petite-Patrie | 1952-1955 | 3 | 3 |
| Lafontaine | AGR5 | Agricultural | Montreal | Le Plateau-Mont-Royal | 1901 | 3 | 1 |
| Lalancette | IND9 | Industrial | Montreal | Mercier-Hochelaga-Maisonneuve | 1922 | 3 | 1 |
| Marguerite-Bourgeoys | AGR6 | Agricultural | Montreal | Le Sud-Ouest | 1913 | 3 | 3 |
| Père-Marquette | IND11 | Industrial | Montreal | Rosemont-La Petite-Patrie | 1953 | 3 | 3 |
| Pointe-Aux-Prairies | IND12, FOR10, AGR11 | Industrial, Forested, Agricultural | Montreal | Rivière-des-Prairies-Pointe-aux-Trembles | 1992 | 8 | 7 |
| Promenade-Bellerive | IND13 | Industrial | Montreal | Mercier-Hochelaga-Maisonneuve | 1978 | 3 | 2 |
| Saint-Gabriel | AGR10 | Agricultural | Montreal | Le Sud-Ouest | 1887 | 2 | 1 |
| Thomas-Chapais | FOR11 | Forested | Montreal | Mercier-Hochelaga-Maisonneuve | 1960 | 3 | 1 |

*(continued)*

| Park | Park.ID | Legacy.ies. | Municipality | Borough | Year Established | Sensors Deployed | Sensors Retrieved |
|---|---|---|---|---|---|---|---|
| Coulée-Grou + Pointe-Aux-Prairies | CON-PaP-CG | Control | Montreal | Rivière-des-Prairies-Pointe-aux-Trembles | | 1 | 1 |
| Promenade-Bellerive | CON-P-B | Control | | | | 1 | 1 |
| Félix-Leclerc | CON-F-L | Control | | | | 1 | 1 |
| Frédéric-Back + Boisé-du-Saint-Sulpice | CON-FB-SS | Control | | | | 1 | 1 |
| Arborétum du Jardin botanique + Lafond + Lalancette | CON-LAL-LAF-JAR | Control | | | | 1 | 1 |
| Lafontaine + Baldwin + Jean-Drapeau | CON-LAF-BAL | Control | | | | 1 | 0 (supplemented sensor) |
| Père-Marquette | CON-P-M | Control | | | | 1 | 1 |
| Marguerite-Bourgeoys + Saint-Gabriel | CON-MB-SG | Control | | | | 1 | 1 |
| Adrien-D-Archambault | CON-ADA | Control | | | | 1 | 1 |
| Arthur-Therrien | CON-AT | Control | | | | 1 | 1 |
| Angrignon | CON-ANG | Control | | | | 1 | 1 |
| Benny | CON-BEN | Control | | | | 1 | 1 |
| Bois-de-Liesse | CON-BDL | Control | | | | 1 | 1 |

*(continued)*

| Park | Park.ID | Legacy.ies. | Municipality | Borough | Year Established | Sensors Deployed | Sensors Retrieved |
|---|---|---|---|---|---|---|---|
| Bois-de-Saraguay | CON-BDS | Control | | | | 1 | 1 |
| Île-Bizard | CON-IB | Control | | | | 1 | 1 |
| Centennial Hall | CON-CENTENNIAL | Control | | | | 1 | 1 |
| Fritz | CON-Fritz | Control | | | | 1 | 1 |
| Cap-Saint-Jacques + L'Anse-à-L'Orme | CON-LALO-CSJ | Control | | | | 1 | 1 |
